# Single cell profiling of Hofbauer cells and fetal brain microglia reveals shared programs and functions

**DOI:** 10.1101/2021.12.03.471177

**Authors:** Alexis M Ceasrine, Rebecca Batorsky, Lydia L. Shook, Sezen Kislal, Evan A. Bordt, Benjamin A. Devlin, Roy H. Perlis, Donna K. Slonim, Staci D. Bilbo, Andrea G. Edlow

**Affiliations:** Department of Psychology and Neuroscience, Duke University, Durham, NC, USA; Research Technology, Tufts Technology Services, Tufts University, Medford, Massachusetts, USA; Division of Maternal-Fetal Medicine, Department of Ob/Gyn, Massachusetts General Hospital, Harvard Medical School, Boston, Massachusetts, USA; Vincent Center for Reproductive Biology, Massachusetts General Hospital Research Institute, Massachusetts General Hospital, Boston, Massachusetts, USA; Department of Pediatrics, Lurie Center for Autism, Massachusetts General Hospital, Harvard Medical School, Boston; Department of Psychiatry and Center for Genomic Medicine, Massachusetts General Hospital, Harvard Medical School, Boston; Department of Computer Science, Tufts University, Medford, MA; Department of Neurobiology, Duke University, Durham, NC 27710, USA; Lurie Center for Autism, Massachusetts General Hospital, Boston, MA

**Author notes:** These authors contributed equally (Alexis M Ceasrine, Rebecca Batorsky, Staci D. Bilbo, Andrea G. Edlow). Correspondence: Andrea G. Edlow, Massachusetts General Hospital, Vincent Center for Reproductive Biology 55, Fruit Street, Thier Research Building, 9^th^ floor, Boston, MA 02114.

**Keywords:** Single-cell RNA sequencing, Hofbauer cells, microglia, sex differences, fetal brain, placenta, development, neuroimmune

## Abstract

Maternal immune activation is associated with adverse offspring neurodevelopmental outcomes, many of which are mediated by *in utero* microglial programming. Microglia remain inaccessible at birth and throughout development, thus identification of noninvasive biomarkers that can reflect fetal brain microglial programming may permit screening and intervention during critical developmental windows. Here we used lineage tracing to demonstrate the shared ontogeny between fetal brain macrophages (microglia) and fetal placental macrophages (Hofbauer cells). Single-cell RNA sequencing of murine fetal brain and placental macrophages demonstrated shared transcriptional programs. Comparison with human datasets demonstrated that placental resident macrophage signatures are highly conserved between mice and humans. Single-cell RNA-seq identified sex differences in fetal microglial and Hofbauer cell programs, and robust differences between placenta-associated maternal macrophage/monocyte (PAMM) populations in the context of a male versus a female fetus. We propose that Hofbauer cells, which are easily accessible at birth, provide novel insights into fetal brain microglial programs, potentially facilitating the early identification of offspring most vulnerable to neurodevelopmental disorders.

## Introduction

Microglia, the resident macrophages of the brain, play a key role in neurodevelopment by modulating synaptic pruning, neurogenesis, phagocytosis of apoptotic cells, and regulation of synaptic plasticity (Bilimoria and Stevens, 2015; Paolicelli et al., 2011; Schafer et al., 2012; Sierra et al., 2010). Aberrant programming of fetal microglia in the setting of maternal immune activation has been identified as a key mechanism underlying abnormal brain development (Fernández de Cossío et al., 2017; Nakai et al., 2003; Nuñez et al., 2003; Smolders et al., 2018; Zhao et al., 2019), and likely contributes to the pathogenesis of neurodevelopmental and psychiatric disorders such as autism spectrum disorder, attention deficit hyperactivity disorder, bipolar disorder, obsessive compulsive disorder and schizophrenia, among others (Bachiller et al., 2018; Frick and Pittenger, 2016; Tay et al., 2018; Williamson et al., 2011). In part because microglia remain inaccessible in fetal life and postnatally, there is currently no way to identify which offspring may be most at risk for adverse neurodevelopmental and psychiatric morbidity after *in utero* exposures. Methods of determining whether or how *in utero* exposures may have primed fetal microglia are urgently needed, to facilitate intervention during critical developmental windows when outcomes can potentially be modified.

Precursors of many tissue-resident macrophages, including microglia, originate in the fetal yolk sac (Ginhoux et al., 2010; Gomez Perdiguero et al., 2015; Pinhal-Enfield et al., 2012; Stremmel et al., 2018). As a pre-placental structure, yolk-sac-derived macrophages likely also colonize the placenta where they are called Hofbauer cells, but prior lineage-tracing has not explicitly focused on the placenta as a resident tissue of interest. Hofbauer cells share exposure to the same intrauterine environment as microglia, and their connection to the developing brain has been posited in their role in the maternal-to-fetal transmission of neurotropic viruses such as Zika, cytomegalovirus, and HIV (Johnson et al., 2018; Jurado et al.; Maciejewski et al., 1993; Noronha et al., 2016; Quicke et al., 2016; Rosenberg et al., 2017; Semmes and Coyne, 2022; Simoni et al., 2017; Zulu et al., 2019). Prior work by our groups has demonstrated that maternal immune activation in the setting of high-fat diet primes placental macrophages and fetal brain microglia in parallel toward a highly-correlated, pro-inflammatory phenotype (Ceasrine et al., 2021; Edlow et al., 2018). Still, whether Hofbauer cells reflect the same transcriptional programs as fetal microglia has not yet been investigated.

To address these gaps in understanding, we sought to definitively trace Hofbauer cells from the fetal yolk sac to the placenta, and to evaluate whether Hofbauer cell programs mirror fetal brain microglial programs, using single-cell RNA-sequencing (scRNA-Seq). Using an inducible macrophage reporter mouse model, we demonstrate that yolk sac-derived macrophages comprise the majority of tissue resident macrophages in both placenta and brain in late embryonic development (embryonic day 17.5; e17.5). Further, we isolated macrophages from wild-type (*C57BL/6J*) placental and fetal forebrain tissues at e17.5 and performed scRNA-Seq, identifying subpopulations of fetal placental macrophages that transcriptionally mirror fetal brain macrophages. While placental macrophage programs mirrored fetal brain microglial programs in both sexes, fetal sex was associated with differential expression of 401 autosomal genes across brain and placental macrophage transcriptomes. Functional analyses of differentially expressed genes provided new insights into sex differences in immune function of both placental and fetal brain macrophages. Taken together, these data suggest that late-term placental macrophages reflect the programs of fetal brain microglia. Hofbauer cells therefore have the potential to provide novel insights into the relationship between placental and brain immune activity and may help identify those offspring most vulnerable to neurodevelopmental morbidity after specific *in utero* exposures.

## Results

### Tissue-resident placental macrophages are yolk sac derived

In both mice and humans, the maternal-fetal interface is comprised of the maternally-derived decidua and the fetally-derived placenta. Fetal placental macrophages have been identified as early as 18 days post conception in humans, and 10 days post conception in mice (e10) prior to the full vascularization of the placenta (Takahashi et al., 1991). This suggests that extra-embryonic placental macrophages (Hofbauer cells), likely derive from the yolk sac, as do many other tissue-resident macrophages, including microglia (Ginhoux et al., 2010).

To determine whether fetal placental macrophages are yolk sac derived, we crossed transgenic mice carrying a floxed *Rosa-tdTomato* allele (Madisen et al., 2010) with inducible transgenic *Csf1R-Cre^ER^* mice to permanently label yolk sac progenitors. Csf1r is active in yolk sac progenitor cells at gestational day 8-9 (gd8-9), thus a 4hydroxytamoxifen (4-OHT) pulse at gd8.5 will label yolk sac-derived macrophages prior to their migration out of the yolk sac (Sasmono et al., 2003). Migration of macrophages out of the yolk sac to colonize the fetal brain and other tissues begins around e9 (Ginhoux et al., 2010), and Csf1r+ placental macrophages are detectable beginning at e10 (Takahashi et al., 1991). Thus, we delivered 4-OHT to Cre^ER^ negative dams (tdTomf/+ or tdTomf/f), crossed with Cre^ER^ positive sires, so that 4-OHT only induced *Csf1R-Cre^ER^* activity in *fetal* cells within the embryos (Figure 1A). We then assessed tdTomato+ cells at e17.5 in placenta and brain from Cre^ER^-negative and Cre^ER^-positive embryos in conjunction with immunohistochemistry for ionized calcium binding adaptor molecule (Iba1), a marker for macrophages. We saw robust colocalization of tdTomato and Iba1 at e17.5 in both the placenta labyrinth (Figure 1B, C’), and fetal brain (Figure 1B, D’, hippocampus shown. We did not see any tdTomato signal in CreER-negative placenta or hippocampus (Figure 1 C, D). Together, these data suggest that resident placental macrophages are primarily derived from yolk sac progenitor cells, similar to fetal microglia.

**Figure 1.**
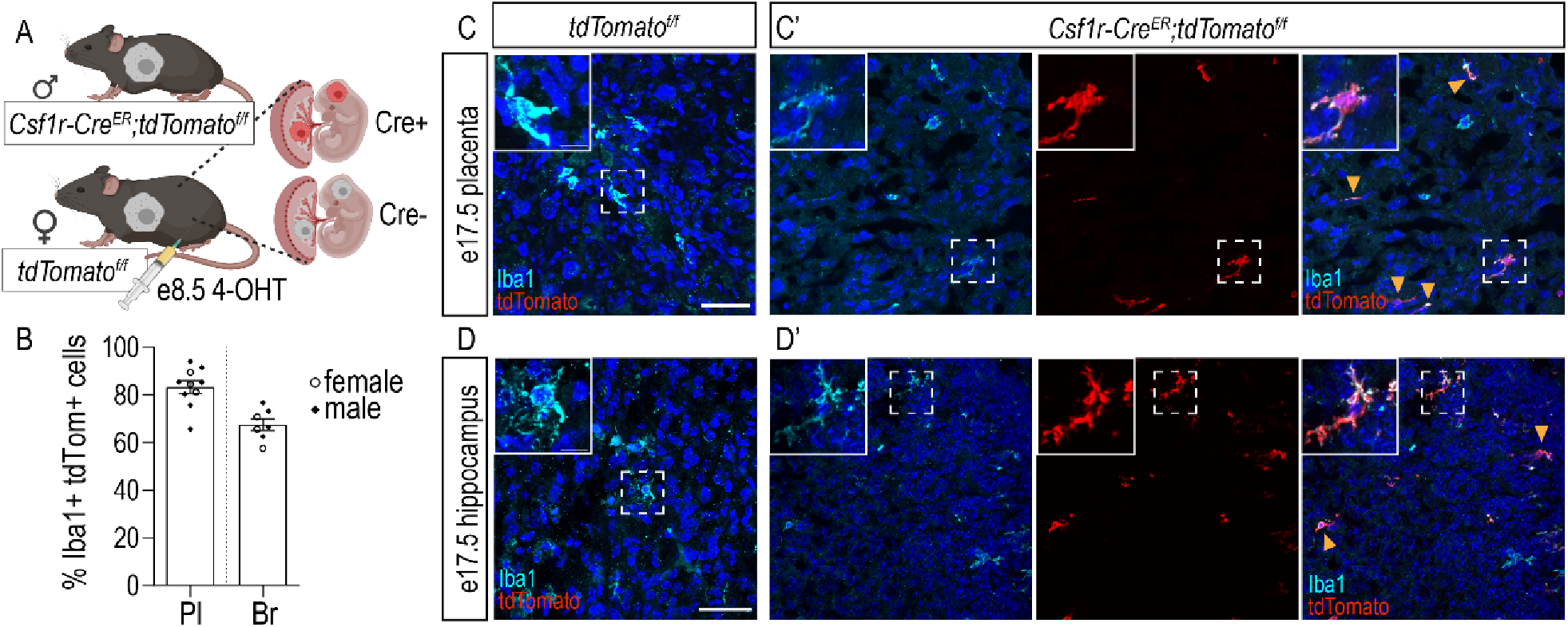
Placental macrophages are yolk sac derived. **A.** Schematic of fetal macrophage labeling. Briefly, male *Csf1R-Cre^ER^;tdTomato^f/f^* mice were timed-mated to female *tdTomato^f/f^* mice. Pregnant females were injected with 4-hydroxytamoxifen (4-OHT) at gestational day 8.5. Embryos were collected at embryonic day 17.5. **B.** Percent of macrophages (Iba1+ cells) labeled with tdTomato in embryonic placenta and hippocampus following 4-OHT administration at e8.5. Open circles represent individual female embryos and closed diamonds represent individual male embryos (n=4 litters). Pl = placenta; Br = brain (hippocampus) **C-C’.** Representative images of Iba1 and tdTomato in control (C) and reporter (C’) placenta from e17.5 embryos. **D-D’.** Representative images of Iba1 and tdTomato in control (D) and reporter (D’) hippocampus from e17.5 embryos. Arrowheads indicate doublepositive (Iba1+ tdTomato+) macrophages/microglia in reporter tissue. Scale 50μm, inset scale 10μm.

### Fetal placental and brain macrophages are heterogeneous populations with shared cluster-specific signatures

To precisely identify subpopulations of fetal placental and brain macrophages, we performed 10X Genomics single-cell RNA-sequencing on macrophage-enriched singlecell suspensions from matched placenta and fetal forebrain tissue from both male and female embryos at e17.5 (Figure S1A). 105,000 cells were sequenced to an average depth of 22,500 reads/cell. These cells represented macrophages and monocytes from matched fetal brains and placentas of 5 females and 4 males from different litters (3 sibling pairs), with an average of 5,886 cells/sample. In order to identify cell types present in our data, we used graph-based clustering followed by identification of cluster-specific marker genes and comparison of marker gene expression and average-expression profiles of all clusters with published placental (Liu et al., 2018; Lu-Culligan et al., 2021; Suryawanshi et al., 2018; Vento-Tormo et al., 2018; Zhou et al., 2021) and fetal brain (Hammond et al., 2019; Kracht et al., 2020; Matcovitch-Natan et al., 2016) datasets (see Methods, Supplemental Figure 1B/C). We selected clusters of macrophage and monocyte cells based on similarity to previously identified gene expression profiles and marker genes, and merged clusters as described in Methods. We included monocyte-like cells in our clustering given recent evidence of potential monocyte-to-macrophage transitional populations at the maternal-fetal interface (Thomas et al., 2020). The final clusters identified are visualized as uniform manifold approximation and projection (UMAP) plots for brain (Figure 2A) and placenta (Figure 2B) along with expression dot plots demonstrating the top three marker genes per cluster. Unless otherwise specified, cluster names begin with the cell type (Mg: microglia, HBC: Hofbauer cell, Mono_FBr: fetal brain monocytes; Mono_FPl: fetal placental monocytes; PAMM: placenta-associated maternal macrophages and monocytes) and end with _top/most significant marker gene in the cluster. Clusters primarily engaged in cell cycle functions (e.g. processes integral to DNA replication) end in _cell cycle. A complete list of cluster marker genes can be found in Supplemental Table 1.

**Figure 2.**
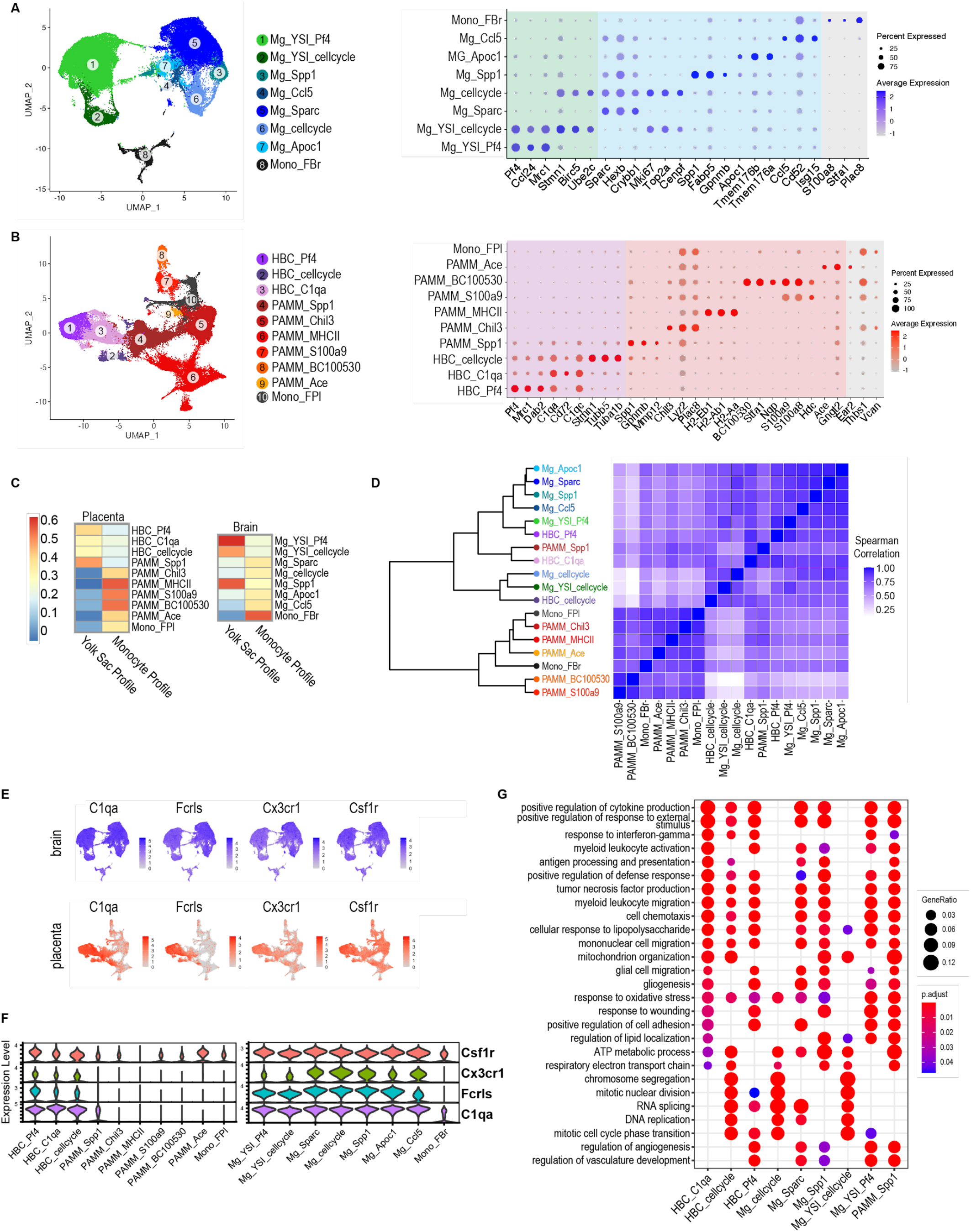
Fetal placental and brain macrophages are heterogeneous populations with shared cluster-specific signatures. **A (left).** Uniform Manifold Approximation and Projection (UMAP) visualization of Cd11b+ macrophageenriched fetal brain microglia/monocyte cells reveals 8 distinct clusters. Unless otherwise specified, clusters are named as “cell type prefix_top marker gene”. Mg: microglia; Mono_FBr: fetal brain monocytes; YSI: yolk sac imprint. **A (right).** Dot plot displaying expression of the top 3 marker genes for each cluster (right), circle size indicates the percent of cells expressing the given gene, color intensity represents the scaled average gene expression. **B (left).** UMAP visualization of Cd11b+ macrophage-enriched placenta macrophage/monocyte populations reveals 4 fetal and 6 maternal clusters. Of note, PAMM_BC100530 was designated a PAMM cluster due to significantly higher expression of *Xist* in a majority of cells compared to *Eif2s3y* and *Ddx3y*, but the presence of Y chromosome marker expression does indicate that this PAMM cluster likely contains some fetally-derived cells (Supplemental Figure 1D). Unless otherwise specified, clusters are named as “cell type prefix_top marker gene”. HBC: Hofbauer cell; PAMM: placenta-associated maternal monocyte/macrophages; Mono_FPl: fetal placental monocytes. Color scheme indicates cell origin (purple=fetal; red=maternal; gray=fetal monocytes). **B (right).** Dot plot displaying expression of the top 3 marker genes for each cluster. Circle size indicates the percent of cells expressing the given gene, color intensity represents the average gene expression. **C.** Heatmap displaying signature enrichment score for yolk sac and monocyte signatures described by Bian and colleagues (Bian et al., 2020; Thomas et al., 2020) **D.** Spearman correlation coefficients between brain and placenta clusters using cluster-average expression of all significant marker genes. Similarity of correlation coefficients across all clusters is displayed as a hierarchical clustering dendrogram. **E/F.** UMAP and Violin plots depicting expression levels of canonical microglia and Hofbauer cell marker genes. **G.** Gene Ontology (GO) Biological Process enrichment analysis for select Microglia, HBC and PAMM clusters. The GO terms displayed here were curated from among the top 25 most significant GO terms, selecting the processes most relevant to macrophage function, and reducing redundancy. Gene Ratio gives the ratio of the number of genes in the query set that are annotated by the relevant GO category and the number of genes in the query set that are annotated in the database of all GO categories. GO terms with an adjusted p-value < 0.05 were considered significantly enriched.

Interrogation of the transcriptional signatures of brain clusters revealed a strong yolk sac signature in three clusters (defined based on the signature of yolk-sac derived myeloid-biased progenitors detailed in Thomas et al., 2020 and Bian et al., 2020; Figure 2C). The clusters with a strong “yolk sac imprint” or YSI signature are named Mg_YSl_cellcycle, Mg_YSl_Pf4, and Mg_Spp1. After distinguishing fetally-from maternally-derived clusters (using Y- and X-chromosome specific gene expression as described below), we identified three fetal placental macrophage clusters with transcriptional profiles most closely related to microglia, including HBC_C1qa, HBC_Pf4, and HBC_cellcycle (Figure 2B–2D). These clusters had marker genes similar to Hofbauer cell clusters described in recent data sets (Lu-Culligan et al., 2021; Suryawanshi et al., 2018; Thomas et al., 2020; Vento-Tormo et al., 2018). We also identified one fetal placental monocyte cluster (Mono_FPl).

Macrophages at the maternal-fetal interface are known to be a heterogenous population consisting of both maternally and fetally-derived cells. While maternallyderived macrophages at the maternal-fetal interface have classically been referred to as decidual macrophages (Mezouar et al., 2021; Mor and Abrahams, 2003; Vento-Tormo et al., 2018), more granular characterization of heterogenous maternally-derived macrophage and monocyte populations has recently been performed (Thomas et al., 2020), with new nomenclature proposed and adopted: placenta-associated maternal monocyte/macrophages (PAMMs) (Semmes and Coyne, 2022; Thomas et al., 2020). Distinct subsets of PAMMs have been identified, including a population of what were previously considered decidual macrophages, and the new identification of separate subtypes of maternally-derived resident placental macrophages distinct from circulating maternal cells (Thomas et al., 2020).

To delineate fetal from maternal macrophages, we evaluated expression of malespecific markers DEAD-Box Helicase 3 Y-Linked (*Ddx3y*) and Eukaryotic translation initiation factor 2 subunit 3, Y-linked (*Eif2s3y*) and female-specific marker X-inactive specific transcript (*Xist*) in brain and placenta clusters from male embryos. *Ddx3y, Eif2s3y*, and *Xist* were selected for their representative expression after interrogation of a broad X- and Y-chromosome-specific gene expression panel, including the antisense *Xist* transcript X (inactive)-specific transcript, opposite strand (*Tsix*), Ubiquitously Transcribed Tetratricopeptide Repeat Containing, Y-Linked (*Uty*), and lysine demethylase 5D (*Kdm5d)* (Supplemental Figure 1D/E). Maternally-derived placental macrophage/monocyte clusters were confirmed via high expression of *Xist* and low/no expression of *Ddx3y* and *Eif2s3y* (Supplemental Figure 1D/E), based on expression in cells isolated from placentas of a male fetus. The same approach was used to ensure isolated brain microglia were fetally-derived (Supplemental Figure 1D/E). Using this approach, we retained 6 PAMM (maternally-derived macrophage) clusters, whose expression profile was similar to recently identified PAMMs from human placenta (Thomas et al., 2020). Via a correlation analysis of overall gene expression across all clusters (see Methods), we identified close relationships between subsets of brain and placental macrophages (Figure 2D). Two major groupings of clusters are noted, one containing Hofbauer cells, microglia, and PAMM_Spp1, and the other containing the rest of the PAMM and monocyte clusters. Not surprisingly given the demonstrated common yolk sac origin, the Hofbauer cell clusters had transcriptional profiles most similar to the “yolk-sac imprint” macrophages Mg_YSI_cellcycle and Mg_YSI_Pf4 (Figure 2D dendrogram and Spearman’s correlation heatmap), and all fetal brain microglial clusters were more closely related to Hofbauer cells than they were to PAMMs, with the exception of PAMM_Spp1. PAMM_Spp1 closely clustered with the largest HBC cluster, HBC_C1qa, suggesting that some immune functions at the maternal-fetal interface may be shared between, or involve close crosstalk between, maternal and fetal cells. In sum, both interrogation of the top marker genes by cluster and correlation analysis of average expression of cluster marker genes demonstrated the closest relationships between Hofbauer cells and fetal brain microglia, particularly a subset of fetal brain microglia with a strong yolk sac-like signature.

Both microglia and Hofbauer cells expressed high levels of canonical microglia/resident tissue macrophage markers *Cx3cr1, Csf1r, Fcrls*, and *C1qa* (Figure 2E/F) among others (Supplemental Figure 1F). The violin plots (Figure 2F) depict expression level of canonical microglial markers in both microglial and placental macrophage clusters depicted. Gene Ontology (GO) Biological Process enrichment analyses of cluster marker genes (Figure 2G) suggest highly conserved biological processes between Hofbauer cells and microglia, and also between Hofbauer cells and PAMM cluster PAMM_Spp1, the PAMM cluster with the strongest yolk-sac signature. While there is some overlap in the processes enriched in all three Hofbauer cell clusters, the largest HBC cluster C1qa is primarily engaged in immune and inflammatory functions (response to interferon gamma, leukocyte activation, positive regulation of cytokine production, and defense response to viruses and bacteria), processes mirrored by microglial clusters Mg_Sparc, Mg_Spp1, and Mg_YSI_Pf4. These same three microglial clusters (Mg_Sparc, Mg_Spp1, and Mg_YSI_Pf4) also share processes in common with the HBC_Pf4 cluster, related to regulation of cell adhesion, angiogenesis and vascular development. HBC_cellcycle cluster is enriched for cell cycle processes such as DNA replication and mRNA processing, functions mirrored by Mg_cellcycle and Mg_YSI_cellcycle. Of note, HBC_cellcycle and Mg_YSI_cellcycle transcriptional signatures may reflect the local proliferation of yolk sac-derived progenitors after trafficking to their final tissue-specific location, as has been previously demonstrated (Thomas et al., 2020). A complete list of enriched GO biological processes in cluster marker genes may be found in Supplemental Table 2.

Shared functions across populations of Hofbauer cells and microglia can be conceptually grouped into five broad categories (Supplemental Figure 2A-E): (1) Cell cycle regulatory functions, including chromosome segregation, mitotic nuclear division, RNA splicing, and mRNA processing (Fig S2A); (2) Immune responses and inflammatory functions, with response to non-self entities a theme repeated across numerous GO terms in both microglia and Hofbauer cell clusters (Fig S2B); (3) Mitochondrial ATP metabolism and energy utilization functions, such as ATP metabolism, mitochondrion organization, and oxidative phosphorylation (Fig S2C); (4) cell migration and cell-cell signaling functions, such as myeloid, leukocyte or macrophage chemotaxis and/or migration (Fig S2D); and (5) lipid transport and localization functions (Fig S2E). In sum, these analyses demonstrate that Hofbauer cells mirror microglia in both gene expression and putative function, supporting the ability of Hofbauer cells to serve as a proxy cell type for microglia at the end of gestation.

#### Similarities between murine and human placental immune single cell gene expression

While mouse microglia are widely accepted as an excellent model for human microglia (Abels et al., 2021; Masuda et al., 2019; Stremmel et al., 2018), less is known about how well mouse placental macrophages model those in humans. Mice and humans are known to share key similarities in immune system development, including the yolk sac as the initial site of hematopoiesis and origin for granulo-macrophage progenitors (such as the CX3CR1+ cells that give rise to microglia and Hofbauer cells) (CindrovaDavies et al., 2017; Godin and Cumano, 2002; Migliaccio et al., 1986; Palis and Yoder, 2001; Takashina, 1987), and conserved cell surface antigen expression in yolk sacderived macrophages (Cindrova-Davies et al., 2017; Stremmel et al., 2018). In addition, the presence of resident fetal macrophages in the mouse placenta is well-documented by our group and others (Chang et al., 1993; Edlow et al., 2018; Gekas et al., 2010; Khalili et al., 1997; Rhodes et al., 2008; Takahashi et al., 1991; Yellon et al., 2019). Despite this knowledge, mouse Hofbauer cells have never been profiled using scRNA-seq, and these data have never been compared to those from human placentas. We therefore compared our mouse data to several human datasets that contained high representation of Hofbauer cells, representing human placentas obtained from 6 weeks through 40 weeks (full term) (Lu-Culligan et al., 2021; Suryawanshi et al., 2018; Vento-Tormo et al., 2018). Comparison of our scRNA-seq data to scRNA-seq data from human placenta datasets demonstrated strong similarities of mouse Hofbauer cells to Hofbauer cell clusters in human placentas at 6-11 weeks (Suryawanshi et al., 2018; Figure 3A/B), 6-14 weeks (Vento-Tormo et al., 2018) Supplemental Figure 3A/B), and full-term (Lu-Culligan et al., 2021) Supplemental Figure 3C/D).

**Figure 3.**
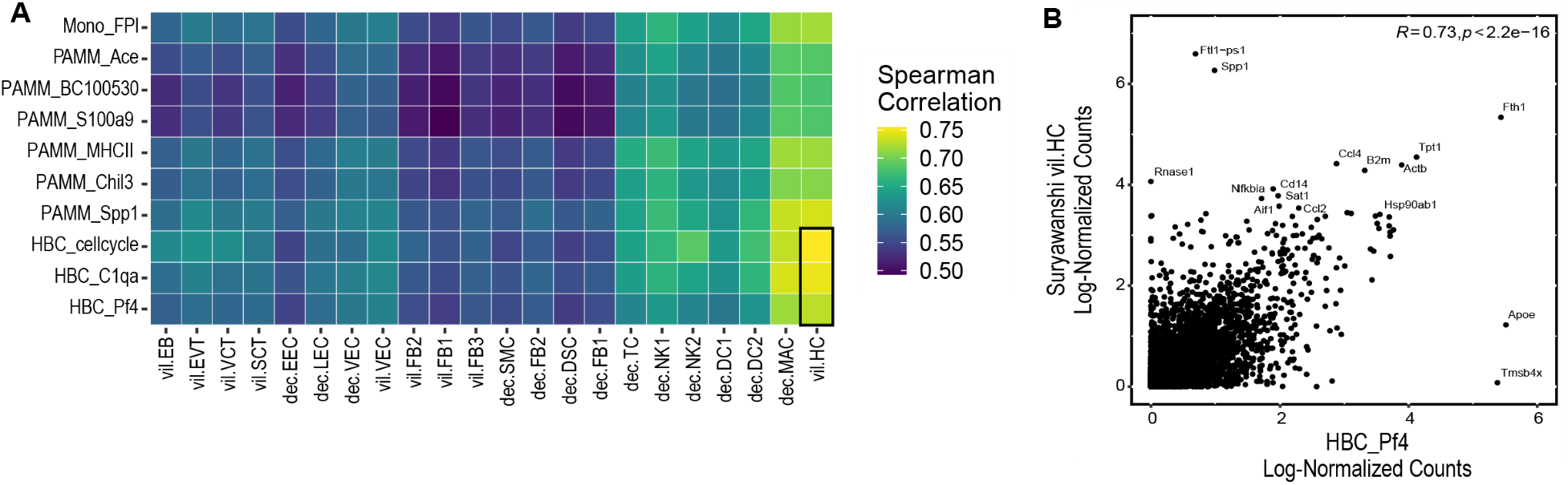
Murine placental macrophages are highly correlated with human placental macrophages. **A.** Heatmap displaying Spearman correlation coefficient R between our murine single-cell placental clusters and previously published human placental clusters (Suryawanshi et al., 2018). Black box outlines correlations between human villous Hofbauer cells (vil. HC, Suryawanshi et al., 2018) and our mouse HBC clusters (HBC_cell cycle, HBC_C1qa and HBC_Pf4). Abbreviations along X-axis of correlation heatmap correspond to single-cell clusters from whole placenta without enrichment for placental macrophages (thus only one Hofbauer cell cluster), and predating PAMM nomenclature. From L to R: Vil: villous, EB: erythroblasts, EVT: extravillous trophoblast, VCT: villous cytotrophoblast, SCT: syncytiotrophoblast, Dec: decidual, EEC: endometrial epithelial cells, LEC: lymphatic endothelial cells, VEC: vascular endothelial cells, FB1-3: fibroblast-like cell clusters 1-3, SMC: smooth muscle cells, DSC: decidualized stromal cells, TC: T cells, NK: natural killer cells, DC: dendritic cells, MAC: macrophages. **B.** Scatter plot demonstrating the significant correlation between our Hofbauer cell cluster most closely related to microglia (HBC_Pf4), and the Hofbauer cell cluster identified by Suryawanshi and colleagues. HBC_Pf4 depicted as representative, but all murine HBC clusters were highly correlated with the human HBC clusters (data in text and Supplemental Figure 3 E/F).

Our murine Hofbauer cell and PAMM clusters mapped closely to human clusters identified as Hofbauer cells (villous Hofbauer cells, vil.HC, HB or vil.Hofb) in these human single cell datasets. Spearman rank correlation demonstrated significant correlation between the transcriptional signature of our murine Hofbauer cell cluster that most closely resembles microglial clusters, HBC_Pf4 (Figure 2D) and human Hofbauer cell signatures in the aforementioned datasets (Figure 3A/B, Spearman’s R = 0.72-0.74, p<2.2e-16, Supplemental Figure 3B,D). In addition to HBC_Pf4, all our murine Hofbauer cell clusters were highly and significantly correlated with human Hofbauer cell clusters, with HBC_C1qa R = 0.75-0.76, p<2.2e-16 (Supplemental Figure 3E) and HBC_cell cycle R = 0.71-0.76, p<2.2e-16 (Supplemental Figure 3F) Taken together, these data demonstrate that placental resident macrophage and monocyte single-cell gene expression signatures are highly conserved across mice and humans.

#### Sex differences in fetal brain and placental macrophage gene expression

Fetal sex is increasingly recognized as a critical factor influencing placental, fetal and neonatal immune responses (Ceasrine et al., 2021; Bordt et al., 2021; Clifton and Murphy, 2004; Edlow et al., 2018; Scott et al., 2009). Sex differences in the incidence and prevalence of many neurodevelopmental and psychiatric disorders are well established, yet the mechanisms underlying sex differences in these disorders are relatively unknown. Given the critical role of microglia in neurodevelopment, sex differences in microglial development and priming are an attractive candidate mechanism underlying sex biases in neurodevelopmental and psychiatric disorders (Lenz and McCarthy, 2015). Understanding how sex contributes to disease development, susceptibility, and severity is of critical importance when considering clinical implications such as treatment efficacy.

To investigate potential baseline sex differences in fetal macrophages, we evaluated differentially expressed genes (DEGs) between males and females by aggregating cells in each individual cluster into a pseudo-bulk count matrix, using DESeq2 to identify DEGs (Methods). Fetal sex did not influence cluster identification or the proportion of cells within clusters (Supplemental Figure 4A/B). Despite the fact that cells were isolated on e17.5, prior to the hormonal surge commonly associated with brain masculinization (Lenz et al., 2013), there were 201 autosomal genes that were differentially regulated in female versus male microglia (Figure 4A/B). This suggests that the local microenvironment or other non-hormonal factors are likely driving differences between male and female microglia in late gestation. There were 27 autosomal genes differentially expressed in female versus male Hofbauer cells and 228 autosomal genes differentially expressed in female versus male PAMMs (Figure 4A/B). Importantly, 17 of the 27 genes that were differentially regulated by fetal sex in Hofbauer cell clusters were also significantly differentially regulated in microglia (Figure 4B), suggesting conserved functionality between the cell types and tissues. Supplemental Table 3 depicts the 17 genes differentially regulated by fetal sex shared between Hofbauer cell clusters and microglia. The direction of dysregulation (e.g. up- versus downregulated) was the same in Hofbauer cells and microglia for all 17 genes.

**Figure 4.**
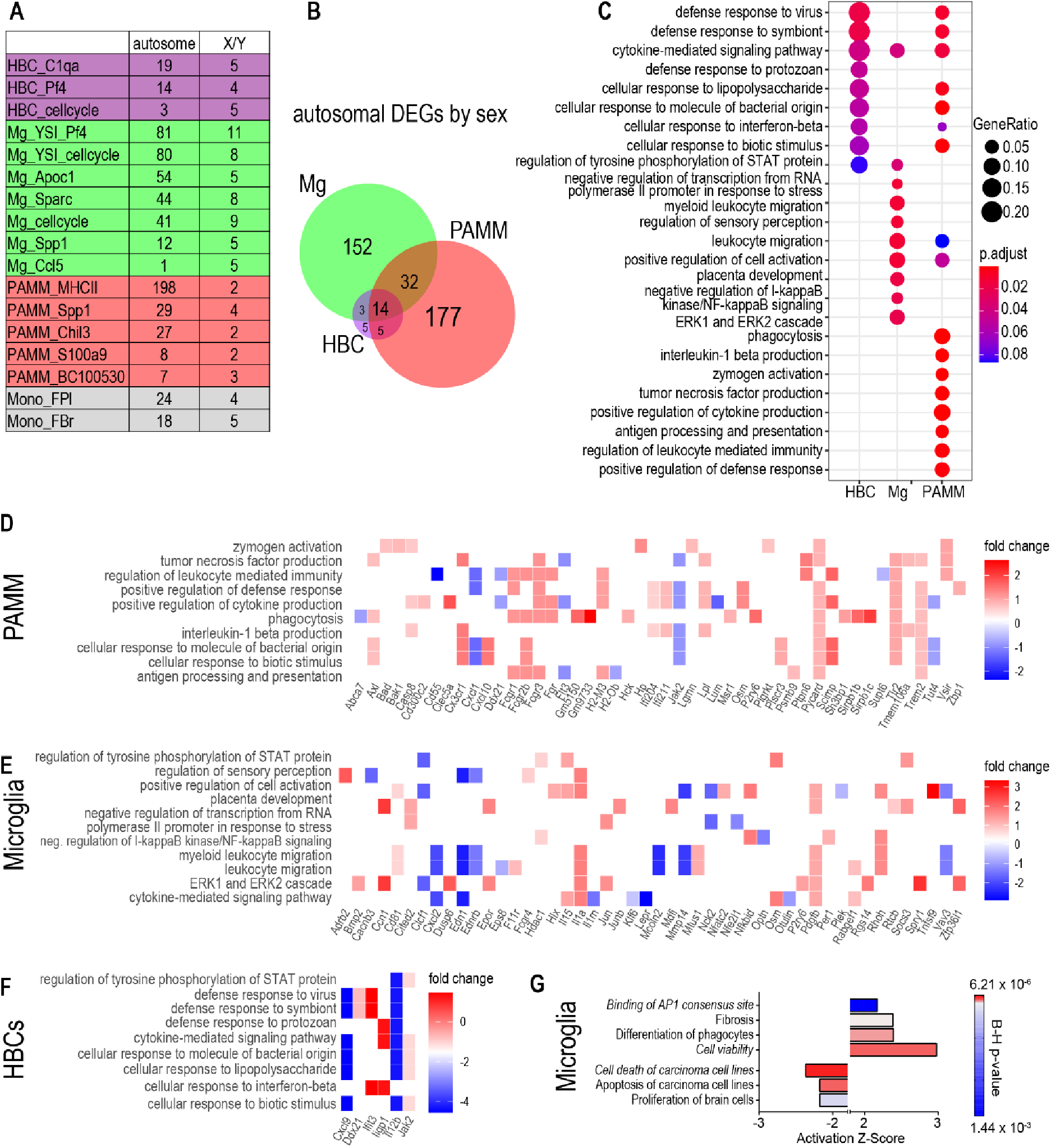
Sex differences in fetal brain and placental macrophage signatures. **A.** The number of differentially-expressed genes (DEGs) between females and males, stratified for genes located on autosomal versus sex chromosomes. **B.** Venn diagram depicting overlap of DEGs by sex grouped by cell type for Mg, HBCs and PAMMs. Note that table values sum to more than the specified number of DEGs between microglia (Mg), Hofbauer cells (HBC) and placenta-associated maternal macrophage/monocytes (PAMM) due to overlap in DEGs across some clusters. **C.** Gene Ontology (GO) Biological Process enrichment analysis for DEGs between males and females grouped by cell type (HBC, Mg, PAMM). The most significant GO terms were curated to select processes most relevant to macrophage function and to reduce redundancy. Gene Ratio gives the ratio of the number of genes in the query set that are annotated by the relevant GO category to the number of genes in the query set that are annotated in the database of all GO categories. GO terms were considered enriched with an adjusted p-value < 0.1. **D-F**. Heat map depicting genes that produce the enrichment for GO terms displayed in E, along with their fold change values. Positive fold change (red) indicated upregulation in females with respect to males in placenta-associated maternal macrophage/monocytes (PAMM) **(D)**, Microglia **(E)** and Hofbauer cells (HBCs, **F**). **G.** Significantly up- and downregulated pathways (absolute Z-score > 2, Benjamini Hochberg p-value < 0.05) identified by Ingenuity Pathways Analysis in female versus male microglia. Terms in italics identify similar pathways implicated in HBCs (IPA could not predict directionality in Hofbauer cells given n=27 DEGs).

##### Functional enrichment analyses of differentially expressed genes by sex

Functional analyses of differentially expressed genes between females and males by cluster was performed using GO Biological Process enrichment in R and Ingenuity Pathways Analysis (IPA). Only differentially expressed autosomal genes were included in the functional analyses. All comparisons are expressed as female versus male, such that “activated” or “inhibited” pathways refers to activated or inhibited in female macrophages relative to male. These analyses suggested significant functional differences in female versus male microglia, Hofbauer cells, and PAMMs in late gestation. Figure 4C depicts differentially regulated biological functions of interest from among the top GO enrichment terms in female versus male microglia, Hofbauer cells, and PAMMs (functional analyses detailed below). Working files for the IPA analyses (Qiagen, Redwood, CA, United States, content version 65367011) may be found in Supplemental Table 4. Significance thresholds within IPA are defined in Methods. Figure 4D-F contain heat maps demonstrating the component genes that produce enrichment for the GO terms displayed in 4C.

##### Shared sex differences in fetal microglial and Hofbauer cell function

Reinforcing the brain-placenta connection, one of the top sex-biased GO biological processes in female versus male microglia was “placenta development”, with this finding primarily driven by the significant upregulation of genes previously implicated in placental extracellular matrix formation, cell migration, RNA repair, cytokine-mediated signaling, monocyte-macrophage differentiation, and angiogenesis (Chen et al., 2015a, 2015b; Imakawa et al., 2016; Ji et al., 2011; Kipkeew et al., 2016; Stumpo et al., 2004) (*Ccn1, Cited2, Epor, Jun, Pdgfb, Rtcb*, and *Zfp36I1*) in female versus male microglia (Figure 4C). Hofbauer cells have been demonstrated to secrete factors that impact placental angiogenesis and remodeling, including IL-8, osteopontin, and matrix metalloproteinase 9 (MMP-9) (Thomas et al., 2020). Our observed pattern of differentially expressed genes in females versus males suggests that female microglia may be more engaged in regulating angiogenesis and vascular remodeling than male microglia (GO analyses suggest that these cells are in clusters Mg_Sparc, Mg_Spp1, and Mg_YSI_Pf4, Figure 2G).

Overall, the GO and IPA analyses suggest a relative upregulation of proinflammatory signaling in female microglia and Hofbauer cells, or a relative repression of proinflammatory signaling in male macrophages. IPA canonical pathway analysis suggested significant activation of multiple pathways related to inflammatory, antiviral and antibacterial signaling in female microglia relative to male, including upregulation of the High Mobility Group-B1 (Z-score 2.0, BH p-value<0.001), IL-1 (Z-score 2.0, BH p-value 0.008), Corticotropin-Releasing Hormone (Z-score 2.0, BH p-value 0.03), Pf3K Signaling in B Lymphocytes (Z-score 1.6, BH p-value 0.003), IL-8 (Z-score 1.3, BH p-value 0.03), and Role of PKR in Interferon Induction and Antiviral Response (Z-score 1.3 BH p-value 0.03) canonical signaling pathways, among others. A complete list of significantly dysregulated canonical pathways in female vs. male microglia may be found in Supplemental Table 5, Tab 1.

Upstream regulator analyses (inferred from DEGs) within IPA also demonstrated significant (BH-p<0.01) upregulation of proinflammatory signaling cascades in female microglia relative to male, including activation of: platelet derived growth factor subunit A and B (*Pdgfa/b*) which regulates IL-6, ERK1/2, FOS, MAPK, and STAT3 signaling; Epidermal growth factor (*Egf*), which regulates ERK 1/2, Akt, MAPK1, STAT3, FOS, PF3K signaling; Fibroblast growth factor 2 (*Fgf2*), regulator of ERK1/2, Akt, MAPK1, FOS, VEGFA, MMP13 signaling; Mitogen activated protein kinase 1 (*Mapk1*) itself; Cd40 ligand (*Cd40lg*), which regulates TNF, IL-1B, IL12B, IL-6, NFKB1 and HLA-DR signaling; and regulators of T-cell gene expression and T-cell receptor signaling such as Nuclear factor of activated T cells 3 (*Nfatc3*). Also activated in female microglia relative to male were upstream regulators involved in regulation of DNA transcription including myocyte enhancer factor 2C and 2D (*Mef2C, Mef2D*), both of which bind multiple histone deacetylases. In addition, numerous apoptosis regulators in the Forkhead family were activated in female versus male microglia, including forkhead box L2 (*FOXL2*), implicated in female sexual differentiation and estrogen receptor binding; forkhead box O1 (*FOXO1*), implicated in cellular glucose homeostasis and response to oxidative stress; and forkhead box O3 (*FOXO3*), a FAS-ligand regulator implicated in apoptosis and autophagy.

Significantly inhibited upstream regulators in female versus male microglia included several genes implicated in epigenetic regulation and transcriptional repression, such as members of the histone deacetylase family (HDACs) and RB transcriptional corepressor 1 (*RB1*), a tumor suppressor implicated in negative regulation of the cell cycle. Additionally, FOS like 1, AP-1 transcription factor subunit (*FOSL1*) signaling, implicated in oxidative stress response was inhibited in female microglia relative to male. These findings suggest that epigenetic regulatory differences may play a role in sexspecific microglial phenotypes. Supplemental Table 5, Tab 2 contains a complete list of significantly activated and inhibited upstream regulators in female versus male microglia.

Supplemental Table 5, Tab 3 describes results of the Downstream Effects analysis within IPA, which evaluates significantly dysregulated cellular and molecular functions and physiological systems in female microglia relative to male, including gene expression patterns consistent with reduced apoptosis, increased cell viability, increased phagocyte differentiation, and increased binding of AP-1 transcription factor to DNA promoter site (Figure 4G).

Sex differences in proinflammatory and immune signaling were also apparent in functional analyses of Hofbauer cell DEGs. GO analyses demonstrated differential responses to interferon-β (IFNβ), defense response to virus, and response to LPS/bacteria, as well as differential signaling through the JAK-STAT pathway in female versus male Hofbauer cells. The top upstream regulators identified by IPA also implicated differential regulation of immune signaling in female versus male Hofbauer cells– pointing to sex differences in Type 1 IFN signaling (IFNα- and IFNβ-mediated signaling), consistent with recently described increased baseline expression of IFN stimulated genes in the Type 1 pathway in female compared to male placentas (Bordt et al., 2021). Other dysregulated upstream regulators in female versus male Hofbauer cells included TANK binding kinase 1 (*Tbk1*), a regulator of interferon regulatory factors IRF3 and 7, and IFNα and -β; pro-inflammatory cytokines including interleukin-4 (*IL-4*), interleukin-6 (*IL-6*), Tumor Necrosis Factor (*Tnf*) and Interferon gamma (*Ifnγ*); regulators of TLR3- and TLR4mediated signaling such as Rho GTPAse activating protein 21 (*Arhgap21*) and *SAM and SH3 domain containing 1* (*Sash1*); and regulators of T-cell receptor signaling and T-cell migration (Dedicator of cytokinesis 8 or *Dock8*, and nuclear factor of activated T cells 2 or *Nfatc2*). This is consistent with prior work on HIV in pregnancy, demonstrating that Hofbauer cells have the ability to engage the T-cell receptor, and can alter T-cell responses and T-cell anergy (Johnson and Chakraborty, 2012). Significantly dysregulated Upstream Regulators, and dysregulated molecular and cellular functions and physiological systems identified by the Downstream Effects analysis in female versus male HBCs are described in Supplemental Table 6, Tabs 1 and 2. In sum, these functional analyses demonstrate significant sex differences in immune function in both microglia and Hofbauer cells, with gene expression patterns suggesting upregulation of inflammatory and immune functions in female cells relative to male, and significant sex differences in apoptosis and autophagy regulation, epigenetic regulation, and vascular remodeling.

##### Fetal sex differences are associated with altered maternal placental macrophage (PAMM) functions

Functional analyses of DEGs in female versus male PAMMs demonstrated significant differences in the maternal immune response to a male versus a female fetus. In the GO Biological Process enrichment analyses, differentially regulated processes in female versus male PAMMs included phagocytosis, bacterial defense response, and proinflammatory cytokine production, including IL-1β (Figure 4C). Although similar upregulation of Type I interferon and other innate immune and pro-inflammatory signaling pathways in female relative to male macrophages were observed in both Hofbauer cells and PAMMs, the number of dysregulated genes by fetal sex and the strength of functional associations were greater for PAMMs than for Hofbauer cells. This suggests that perhaps both fetal and maternal placental resident macrophages have dampened proinflammatory responses in a male pregnancy, with maternal macrophages more significantly impacted, highlighting a potential role for maternal more than fetal placental macrophages in sexspecific immune tolerance of a male versus female fetus.

IPA analyses of sex differences in PAMM functions echoed many of the same themes identified by GO enrichment. Canonical pathways analyses identified significant activation of the “Role of Hypercytokinemia/Hyperchemokinemia in the Pathogenesis of Influenza” canonical pathway (Z-score 2.5, BH p-value 0.03) in female versus male macrophages. This pathway is implicated in antiviral/innate immune response, and involves signaling through the Type I, II and III Interferon pathways, with upregulation of TNF-α, IL-1, IL-6, MIP-1α, MCP-1/CCL, IP-10, and RANTES/CCL5. Upstream regulator analyses (inferred from DEGs) also suggested significant upregulation of Type I, II and III IFN signaling in female relative to male PAMMs, with predicted activation of IFN-α, IFN-β, interferon alpha and beta receptor subunit 1 (*Ifnar1*), IFN lambda 1 (*Ifnl1*), IFN gamma (*Ifnγ*), Tnf superfamily (*Tnf*), nuclear factor kappa B subunit 1 or *Nfκϐ1*, several interferon regulatory factors including *Irf7, Irf1, Irf3, Irf5*, and pro-inflammatory toll-like receptormediated signaling, including activation of Myeloid differentiation primary response gene 88 (*Myd88*), and toll-like receptor 9 (*Tlr9*), among other activated upstream regulators. Inhibited upstream regulators in female relative to male PAMMs included regulators implicated in ATPase binding and NRF2-mediated oxidative stress response (caseinolytic mitochondrial matrix peptidase proteolytic subunit or *Clpp*), in C-C chemokine signaling (atypical chemokine receptor 2 or *Ackr2*), regulation of autophagy and cell death (immunity-related GTPAse family M member 1 *Irgm1*), and epigenetic regulation via chromatin binding (Cbp/p300 interacting transactivator with Glu/Asp rich carboxy-terminal domain 2 or *Cited2*). A complete description of significantly dysregulated Upstream Regulators in female versus male PAMMs can be found in Supplemental Table 7, Tab 1. In the Downstream Effects analysis, numerous molecular and cellular functions were predicted to be significantly increased (activation Z-score ≥2, BH p-value < 0.05) in female relative to male PAMMs, including innate immune responses, phagocytosis, immune cell recruitment, carbohydrate metabolism, and apoptosis and cell death (Supplemental Figure 4C; Supplemental Table 7, Tab 2). Functions predicted to be decreased included multiple terms related to proliferation of lymphocytes and lymphoid cells. Taken together, these functional analyses demonstrate significant sex differences in maternal placental macrophage programs, with relatively reduced immune and inflammatory signaling and reduced phagocytic functions in male PAMMs, suggesting an important role for maternal placental-associated macrophages in immune tolerance of a male fetus.

## Discussion

Microglia, brain resident macrophages, are particularly vulnerable to the *in utero* environment, and diverse maternal immune-activating exposures—including bacterial/ viral infection, metabolic inflammation (e.g. obesity, diabetes), environmental toxicants, and maternal stress—can impact critical microglial functions, including alterations in synaptic pruning or pro-inflammatory cytokine release in response to local stimuli. This programming in turn may contribute to adverse neurodevelopmental and psychiatric outcomes in offspring ranging from autism spectrum disorder/social behavioral deficits, to attention deficit hyperactivity disorder, cognitive deficits, schizophrenia, obsessive compulsive disorder, anxiety, and depression (Bachiller et al., 2018; Bilbo and Tsang, 2010; Bilbo et al., 2005; Chen et al., 2021; Edlow, 2017; Frick and Pittenger, 2016; Hanamsagar and Bilbo, 2017; Nogueira Avelar E Silva et al., 2021; Shook et al., 2020; Tay et al., 2018; Van Lieshout and Voruganti, 2008; Williamson et al., 2011). However, the mechanisms that dictate susceptibility are unclear, and there is a critical need to identify offspring most vulnerable to neurodevelopmental morbidity after potentially harmful *in utero* exposures. We therefore sought to evaluate whether extra-embryonic placental macrophages—Hofbauer cells—could serve as a proxy cell type for fetal brain microglia. Here, we demonstrate (1) a common yolk sac origin for microglia and Hofbauer cells, (2) mirrored transcriptional and functional programs between subsets of microglia and Hofbauer cells, providing strong evidence that late-term Hofbauer cells have the potential to inform on the programs of fetal microglia, (3) significant correlations between murine and human placental macrophage populations, and (4) significant sex differences between female and male microglial and placental macrophage programs, consistent with increased inflammatory and immune tone and phagocytic functions (upregulated baseline state) in female microglia and placental macrophages relative to male.

Fate-mapping studies demonstrate that primitive yolk sac-derived macrophages colonize the developing brain as early as embryonic day 9.5 (e9.5) in rodents, entering the parenchyma via the blood stream and ventricles (Ginhoux and Prinz, 2015; Ginhoux et al., 2010; Gomez Perdiguero et al., 2015; Stremmel et al., 2018; Takahashi et al., 1991). Microglia continue to proliferate throughout the first postnatal weeks in humans and rodents, forming a self-renewing pool that lasts throughout the lifespan, without contribution from peripheral hematopoietic cells under normal conditions (Ajami et al., 2007; Bruttger et al., 2015; Ginhoux and Prinz, 2015; Ginhoux et al., 2010; Gomez Perdiguero et al., 2015; Greter et al., 2015; Salter and Stevens, 2017). Because fetal yolk sac-derived macrophages are the progenitors for the pool of microglia throughout an individual’s lifetime (Ginhoux and Prinz, 2015; Ginhoux et al., 2010; Gomez Perdiguero et al., 2013, 2015), inappropriate fetal macrophage priming (“trained immunity” (Haley et al., 2019; Li et al., 2018)) due to *in utero* immune activation may have lifelong neurodevelopmental consequences. It is therefore critical to identify a non-invasive readout for microglial state at birth, to identify the most vulnerable offspring and facilitate screening and possibly intervention during key developmental windows.

While limited data suggest that Hofbauer cells have strong immune functional correlation with microglia (Edlow et al., 2018; Na et al., 2021), no previous studies have investigated whether murine Hofbauer cells transcriptionally reflect microglia, and whether they can be used to model human Hofbauer cells. Parallels between human and murine placentation are apparent in both gross morphology and function as well as molecular pathways. Although the mouse placenta does have certain fundamental differences from human, such as the degree of contact between fetal tissues and maternal blood, or variability in tissue structure between human placental villi and mouse fetal placental labyrinth, both species share important functions in nutrient transport, blood filtration, and immunocompetency (Hemberger et al., 2020; Moffett and Loke, 2006; PrabhuDas et al., 2015). Key similarities between mouse and human immune system development relevant to these experiments include the yolk sac as the initial site of hematopoiesis and origin for both microglia and Hofbauer cells (Cindrova-Davies et al., 2017; Godin and Cumano, 2002; Migliaccio et al., 1986; Palis and Yoder, 2001; Takashina, 1987), but certainly differences remain between human and mouse immune development (Mestas and Hughes, 2004). To establish the extent to which murine placental macrophages can be used to model human Hofbauer cells and PAMMs, we interrogated the transcriptional relationships between our Hofbauer cell and PAMM clusters and those of three independent published human data sets (Lu-Culligan et al., 2021; Suryawanshi et al., 2018; Vento-Tormo et al., 2018). In all comparisons, the signatures of murine Hofbauer cells and PAMMs from our dataset were closely correlated with human Hofbauer cell and PAMM signatures, suggesting a highly conserved evolutionary relationship. Taken together, these findings provide strong evidence that the mouse placenta can be used to understand immune interactions at the maternal-fetal interface.

Despite their recognized importance in immune signaling at the maternal-fetal interface (Reyes and Golos, 2018), Hofbauer cells remain under-characterized due to challenges in cell isolation and maintenance of viability (Megli and Coyne, 2020). Thus, while microglia have been demonstrated to be a highly heterogeneous cell type with subsets of cells performing specialized functions (Hammond et al., 2019; Kracht et al., 2020), there is a relative lack of knowledge regarding Hofbauer cell subsets and their functions. By enriching for macrophages and monocytes with a Percoll gradient followed by sub-selection for CD11b+ cells, our experiments were able to provide uniquely targeted sequencing and greater resolution on placental macrophage populations than has been previously achieved. This increased resolution led to new transcriptional and functional insights on the under-characterized heterogeneity of Hofbauer cells and PAMMs. Consistent with other groups (Thomas et al., 2020; Tsang et al., 2017; VentoTormo et al., 2018), we found that maternal cells contribute significantly to placental macrophage populations. Our use of male placentas and a panel of X- and Ychromosome markers to identify fetal versus maternal placental macrophages represents an advance in understanding, permitting more precise characterization of the transcriptional profile and function of fetal versus maternal placental macrophages.

Our findings also confirm and expand upon the recent discovery of PAMMs (Thomas et al., 2020), identifying further resolution on subclusters of PAMMs (six versus three clusters), which in turn permitted new insights into their function. Examination of the hierarchical clustering and GO enrichment analyses of the PAMM_Spp1 cluster, a cluster similar to the PAMM1a cluster identified by Thomas and colleagues, provides additional insights into function. Cells in the previously-described PAMM1a cluster were noted to be in direct contact with the (fetal) syncytiotrophoblast layer of the placenta, with these PAMM cells enriched in lipid droplets (Thomas et al., 2020). It was therefore hypothesized that these cells might be important in facilitating the survival, integrity, and repair of the syncytiotrophoblast layer of the placenta. Consistent with this observation of close crosstalk and engagement of these maternal macrophages with fetal placental cells, our PAMM_Spp1 cluster, unlike other PAMM clusters, had substantial overlap in its transcriptional and functional profile with both fetal brain microglia and Hofbauer cells (Figure 2D and G). Also consistent with the finding that PAMMs implicated in repair and regeneration of the fetal cells were localizing to lipid droplets adherent to the syncytiotrophoblasts (Thomas et al., 2020), we found that GO terms related to regulation of lipid localization and cellular response to lipid, and response to wounding also mapped to PAMM_Spp1 (Figure 2G and Supplemental Figure 2E), as well as to two HBC (HBC_C1qa and HBC_Pf4) and three microglial clusters (Mg_Sparc, Mg_Spp1, and Mg_YSI_Pf4). Interrogating the function of microglia in these clusters will be an interesting direction for future work. It has been demonstrated that microglia can accumulate lipid droplets, in some cases leading to impaired phagocytosis and inflammation (Marschallinger et al., 2020). Taken together, these findings demonstrate the potential for scRNA-seq signatures to generate insights into function of subpopulations of Hofbauer cells, microglia, and maternally-derived placental macrophages, as well as the potential for Hofbauer cells to mirror microglia in both gene expression profile and function.

Prior work has demonstrated sex-specific fetal brain gene expression signatures using whole fetal forebrain in a murine model, without subselection for any specific cell or immune cell subtype (Edlow et al., 2016b). Given a strong sex bias in many microglial mediated neurodevelopmental disorders, with autism spectrum disorder, attention deficit hyperactivity disorder, and cognitive delay/learning disabilities all more common in males than females (Bordeleau et al., 2019; Hanamsagar and Bilbo, 2016; Lenz and McCarthy, 2015; Shook et al., 2020), and the fact that male fetal macrophages (both brain and placental) may be more vulnerable to pro-inflammatory intrauterine priming compared to female macrophages (Edlow et al., 2018; Na et al., 2021), we sought to investigate potential baseline sex differences in fetal macrophage populations. Immune responses at the maternal-fetal interface are influenced by both maternally- and fetally-derived macrophages. By excluding X and Y-chromosome genes from our sex differential gene expression analyses, we were able to gain greater resolution on autosomal genes and functions that differ by sex, compared to prior studies (Cvitic et al., 2013; Gonzalez et al., 2018; Sun et al., 2020). Our finding that female placental and brain macrophages have relatively increased immune/inflammatory “tone” relative to male are consistent with other groups who have demonstrated upregulation of innate-immune-regulating genes in female versus male placental cells at baseline (Bordt et al., 2021; Kieffer et al., 2018; Rosenfeld, 2015; Sun et al., 2020); taken together, these findings suggest that female fetuses may be better equipped to respond to immune challenge during key developmental windows. Fetal sex exerted a greater influence on gene expression in maternally-derived macrophage clusters (PAMMs) compared to fetal Hofbauer cells, with nearly 10 times more DEGs identified in PAMM versus Hofbauer cell clusters. GO analysis revealed a striking fetal sex-dependent response in immune-related PAMM functions such as phagocytosis, response to bacterial and viral origin, and interleukin production, suggesting that maternally-derived placental macrophage populations play an important role in immune tolerance of a male versus female fetus. In sum, our analyses of sex differences in PAMMs suggest that the maternal immune system is finely-tuned to locally respond to a male or female fetus, with crosstalk between subsets of fetal and maternal placental macrophages.

Our study has potential limitations. Methods used to digest brain tissue and isolate microglia vary widely but can substantially alter the transcriptomic and functional capabilities of these cells. Some have asserted that positive cell selection with microbeads may be associated with cell activation (Bhattacharjee et al., 2018), but other methods necessary for cell isolation, such as Fluorescence activated cell sorting or FACS, demonstrate similar induction of markers associated with microglial activation (e.q. CD86 and TLR4) (Ayata et al., 2018; Haimon et al., 2018). In addition, while other studies have utilized different methods for both microglial and placental macrophage cell isolation (Hammond et al., 2019; Thomas et al., 2020), many groups have reported similar nonspecificity of their “Hofbauer cell” isolation protocols that also result in the isolation of maternal macrophages. CD11b+ bead isolation following a 70/30 Percoll gradient was selected due to its ability to permit isolation of fetal brain and placenta macrophages in parallel, in a timely fashion to optimize cell viability for sequencing. Fortunately, singlecell RNA-seq coupled with the use of Y-chromosome specific markers in male embryos permitted resolution of maternal from fetal macrophages. ScRNA-seq also provided the advantage, compared to pre-sorting with FACS followed by bulk RNA-seq, of permitting post-sequencing resolution of contaminating cell types, such as fibroblasts, vascular endothelial cells, and natural killer cell populations. While these investigations were designed to answer the question of whether Hofbauer cell subsets transcriptionally reflect fetal brain microglial signatures, confirmation of the biological functions of highly correlated placental and brain clusters is beyond the scope of this investigation. Further investigation of the putative biological functions of clusters of interest and evaluation of alterations to these baseline signatures in specific maternal exposure and disease states will be important directions for future work.

In summary, these data provide a precedent for using placental Hofbauer cells as a noninvasive surrogate for fetal brain microglial programming. Given the shared transcriptional programs of healthy fetal brain microglia and Hofbauer cells in both males and females, we hypothesize that Hofbauer cells and microglia will respond similarly to maternal exposures. This is supported by previous work from our groups and others demonstrating that Hofbauer cells and fetal brain microglia respond similarly to bacterial infection and to the bacterial endotoxin LPS after priming by maternal obesity (Edlow et al., 2018; Na et al., 2021). This work paves the way for longer-term translational studies in humans correlating Hofbauer cell immune profiles with offspring neurological outcomes. Such studies will lend critical evidence to whether Hofbauer cells can serve as an early indicator of neurodevelopmental risk, with the ultimate goal of identifying vulnerable offspring at birth and facilitating interventions during key developmental windows of plasticity. Exploring how Hofbauer cells respond to a variety of maternal influences will provide further insight into the biology of microglial priming and can inform potential targeted therapeutics.

## Methods

### Resource Availability

#### Lead contact

Further information and requests for resources/reagents should be directed to and will be fulfilled by the lead contact, Andrea Edlow

#### Materials availability

This study did not generate new unique reagents

#### Data and code availability

Single-cell RNA-seq data will be deposited in GEO prior to publication. Accession numbers will be listed in the key resources table. Any additional information required to reanalyze the data reported in this paper will be available from the lead contact upon request.

### Experimental Model and Subject Details

#### Animals

##### Strains and husbandry conditions

All procedures relating to animal care and treatment conformed to Massachusetts General Hospital Center for Comparative Medicine Program, Duke University Animal Care and Use Program, and NIH guidelines. Animals were group housed in a standard 12:12 light-dark cycle. The following mouse lines were used in this study: *FVB-Tg(Csf1rcre/Esr1*)1Jwp/J* (Jackson Laboratory, stock no. 019098, referred to as *Csf1r-Cre^ER^* hereafter), *B6.Cg-Gt(ROSA)26Sor^tm14(CAG-tdTomato)Hze^/J* (Jackson Laboratory, stock no. 007914, referred to as *tdTomato^f/f^* hereafter), and *C57BL/6J* (Jackson Laboratory, stock no. 000664). *Csf1r-Cre^ER^* animals were backcrossed to *C57BL/6J* mice for one generation prior to breeding with *TdTomato^f/f^* animals. *Csf1r-Cre^ER^* was maintained in the males for all experimental studies, and genotyping was performed per Jackson Laboratory published protocols for each strain.

##### Transgenic Breeding/Maintenance

Male *Csf1r-Cre^ER^;TdTomato^f/f^* mice were crossed with *TdTomato^f/f^* or *Tdtomato^f/+^* females to generate control *TdTomato^f/f^* or *Tdtomato^f/+^* and experimental *Csf1r-Cre^ER^;TdTomato^f/f^* or *Csf1r-Cre^ER^;TdTomato^f/+^* animals within the same litters. We did not see any spontaneous recombination (e.g. tdTomato fluorescence or RFP immunoreactivity) in control animals. Pregnancy was determined by the presence of a copulation plug (gestational day 0.5 (gd0.5)), and maternal weight was measured at gd0.5 and gd8.5 (to confirm pregnancy weight gain). Pregnant females were injected intraperitoneally (i.p.) with 10 mg/kg 4-hydroxytamoxifen (4-OHT, Millipore-Sigma cat #H6278) dissolved in corn oil (Sigma) at gd8.5 and euthanized at gd17.5 with CO_2_ followed by rapid decapitation. Sex was recorded and is clearly noted throughout the manuscript.

### Method Details

#### Tissue collection for immunohistochemistry

Uterine horns containing embryos were rapidly dissected and placed on ice in sterile 1X PBS. Individual embryos were separated, and placenta, brain, and tail tissue (for genotyping) were collected. Placenta and brain tissue were fixed in 4% paraformaldehyde in PBS (PFA, Sigma) overnight at 4°C, cryoprotected in 30% sucrose + 0.1% sodium azide in PBS (Sigma), and embedded in OCT (Sakura Finetek, Torrance, CA) before being cryo-sectioned. Sections were frozen at −80°C for storage. 40μm cryosections were collected directly onto Superfrost slides (Fisher), permeabilized in 1% Triton X-100 in PBS, and blocked for 1 hr at room temperature using 5% goat serum (GS) in PBS + 0.1% Tween-20. Sections were then incubated for 2 nights at 4°C with chicken anti-Iba1 (Synaptic Systems, 234 006) and rabbit anti-RFP (Rockland, 600-401-379). Following PBS washes, sections were then incubated with anti-rabbit Alexa-594 (placenta and brain), anti-chicken Alexa-647 (placenta) or anti-chicken Alexa-488 (brain) (1:200; ThermoFisher), and DAPI (100μg/mL). Ten z-stacks of 0.5μm optical thickness were taken using a Zeiss AxioImager.M2 (with ApoTome.2) from at least 5 sections (each 400μm apart) from the fetal compartment of each placenta. Ten z-stacks of 0.5μm optical thickness were also taken from at least 3 hippocampal sections (each section being 200μm apart).

#### Tissue collection for sequencing

Pregnant *C57BL/6J* dams were euthanized at gd17.5. Uterine horns containing embryos were rapidly dissected and embryos were placed on ice in sterile PBS. Brains and placentas were isolated, and fetal forebrain and placenta were diced finely with sterile blade and placed into collagenase A digestion solution (Millipore-Sigma; 11088793001) on ice. Cd11b-positive cells were isolated from both placenta and brain tissue as previously described (Bordt, 2020; Edlow et al., 2018). Briefly, samples were serially dissociated into a single cell suspension using hand-flamed Pasteur pipettes, and the resultant suspension was enriched for macrophages and monocytes using a 70%/30% Percoll gradient (Millipore-Sigma; GE17-0891-01). This enriched macrophage/monocyte solution was incubated with human and mouse CD11b microbeads (Miltenyi Biotec; 130093-634), and cells were further enriched for Cd11b-positive macrophages/monocytes using Miltenyi MACS Separation Columns (LS; 130-042-401) and a Miltenyi QuadroMACS Separator (130-090-976). Fresh CD11b+ cells from 4-5 biological replicates per group (4 male brains and matched placentas, 5 female brains and matched placentas) were then prepared for single-cell RNA sequencing (10X Genomics v3.0 chip). Approximately 5,900 cells/sample were sequenced, with a total of 105,000 sequenced cells analyzed across 18 samples. Data from pilot and full-depth batches were integrated for final analysis (pilot runs averaged 5,810 reads/cell and full-depth 22,563 reads/cell).

### Quantification and Statistical Analysis

#### Immunohistochemistry quantification

Maximum intensity projections were generated with FIJI (FIJI Is Just ImageJ(Schindelin et al., 2012)), and the Cell Counter plugin was used to aid in manual counting of Iba1 + and tdTomato/RFP+ cells. Counts were done by an individual blinded to the age, sex, and genotype of the tissue. Sample sizes can be found in the legend for Figure 1.

#### Single cell RNA-sequencing analysis

10x Data analysis and clustering: 10x scRNA-seq data was aligned with 10x Genomics Cell Ranger (v3.0) (Zheng et al., 2017) against the mm10 Mouse reference genome. Downstream analysis was performed with R (v.4.0.0) package Seurat (v4.0.3) (Butler et al., 2018; Hao et al., 2021; Satija et al., 2015; Stuart et al., 2019). Initial filtering was performed on each sample as follows: cells with UMI count <500, gene count <250, or fraction of reads aligning to mitochondrial genes > 0.2 were removed; genes detected in < 10 cells were also removed. All samples were normalized and putative doublet cells were removed using predictions generated from DoubletFinder (v2.0.3) (McGinnis et al., 2019). All samples were integrated to remove batch effects from individual animals using the Seurat Single Cell Transform workflow (Hafemeister and Satija, 2019) using the top 3000 variable features, followed by clustering using the Leiden algorithm on the shared nearest neighbor graph, as implemented in the Seurat function FindClusters function with resolution 0.4, and visualization by UMAP using the first 50 principal components. Cluster marker genes were identified using Seurat function FindAllMarkers. Markers were considered significant with adjusted p-value < 0.05, expression in >25% of cells in the cluster under consideration, and average log fold-change between cells in and out of the cluster under consideration >0.25. To compare the cluster signatures identified in brain and placenta, starting with the total set of significant marker genes in all clusters (1651 genes), we calculated the average expression profiles of these genes for each cluster, and visualized the correlation of marker-gene average expression profiles between all clusters.

##### Cluster annotation

Identification of UMAP cell clusters with cell types was done in three steps: **(1)** First, clusters were annotated using R package SingleR (v1.2.4) with celldex (v.0.99.1) (Aran et al., 2019) packages built-in MouseRNAseq reference; **(2)** annotation was confirmed using the top Spearman correlation coefficient between cluster-averaged gene expression in our clusters and cluster-averaged gene expression cell types from reference datasets for mouse microglia (Hammond et al., 2019; Kracht et al., 2020) and human placenta (Lu-Culligan et al., 2021; Suryawanshi et al., 2018; Vento-Tormo et al., 2018) using the cluster correlation method recently described (Lu-Culligan et al., 2021) (results shown in Figure 3 and Supplemental Figure 3). Scatterplots and heatmaps were produced using the tidyverse 1.31 R library collection (Wickham et al., 2019); **(3)** celltype assignment was refined using manual examination of marker genes. Clusters identified as Monocyte- or Macrophage-like were selected and re-clustered (the full set of identified clusters is shown in Figure S1B). When comparing the correlation between cluster-averaged gene expression between brain and placenta samples (Figure 1D), we restricted the expression matrix to significant marker genes from brain and placenta clusters. Single-cell gene signature enrichment scores were calculated using the AddModuleScore function in Seurat using gene signatures described by Thomas et al (Thomas et al., 2020).

##### Differential gene expression analysis

Differentially expressed genes (DEGs) between males and females were calculated by aggregating counts by cluster and by individual into a pseudobulk count matrix (as implemented by the Libra R package (Squair et al., 2021)), and using the Wald test for differential expression in DESeq2 (Love et al., 2014). Recent benchmarking studies have demonstrated that bulk RNAseq methods perform equally well to those developed for scRNAseq while also controlling false discovery rates (Soneson and Robinson, 2018; Squair et al., 2021). For our hypothesis-generating analyses (e.g. sex differences in microglia, hofbauer cells, and PAMMs), we considered genes with adjusted p-value < 0.1 to be differentially expressed. Genes whose significance was driven by a single sample were eliminated.

##### Functional gene analyses (GO and IPA)

Gene ontology enrichment analysis was performed on both cluster marker genes and sex-differentially expressed genes using R package clusterProfiler (v. 4.0.5) (Wu et al., 2021) enrichGO function and underlying database AnnotationDb org.Mm.eg.db (v3.13.0). For marker genes, enrichment analysis was performed per cluster and GO terms were considered significant for marker gene enrichment with an adjusted p-value < 0.05. For sex-differentially expressed genes, we grouped DEG by cell type for HBC, Mg and PAMM clusters prior to enrichment analysis and considered GO terms significant with an adjusted p-value less than 0.1. GO terms shown for marker genes in Figure 2G and DEG in Figure 4C were curated to show the most significant terms in selected clusters that were relevant to macrophage function, and to remove redundancy.

For the Downstream Effects analyses, IPA pathways (cellular and molecular functions and physiological systems) were defined as significantly dysregulated if their Zscores were ≥ 2 (activated) or ≤ 2 (inhibited), if they had Benjamini Hochberg-corrected p-values < 0.05, and contained 3 or more genes from our differentially-expressed gene set, consistent with prior work and IPA’s recommended thresholds for significance (Edlow et al., 2016a, 2015, 2016b, 2019; Ingenuity Systems, 2021a). Only Canonical Pathways and Upstream Regulators with predicted activation or inhibition (Z-scores ≥ 2 being activated and ≤2 inhibited) and/or Benjamini-Hochberg p-value < 0.01, reflecting 3 or more genes in our differentially-expressed set were considered significant, consistent with IPA’s recommended thresholds for causal pathway analyses (Ingenuity Systems, 2021b; Krämer et al., 2014).

## Supplemental Files

### Supplemental Figures may be found in Supplemental Figures pdf Supplemental Tables

**Supplemental Table 1.** Significant Cluster Marker Genes

Footer: Tab 1, Brain. Tab 2, Placenta. The following filtering thresholds were applied: adjusted p-value < 0.05, expression in >25% of cells in the cluster under consideration (pct.1>0.25), and average log fold-change between cell inside and outside of the cluster under consideration > 0.25 (avg_Log2FC>0.25). Cluster: name of the cluster under consideration; p_val: p-value unadjusted; avg_log2FC: Log2 fold-change of the average expression between the two groups (inside and outside the cluster under consideration)-positive values indicate that the gene is more highly expressed in the first group; pct.1: The percentage of cells where the gene is detected in the cluster under consideration; pct.2: The percentage of cells where the gene is detected in clusters outside the cluster under consideration; p_val_adj: Adjusted p-value, based on Bonferroni correction using all genes in the dataset.

**Supplemental Table 2.** GO Biological Processes Enriched in Cluster Marker Genes

Footer: GeneRatio: The ratio of the number of genes in the query set that are annotated by the GO ID and the number of genes in the query set that are annotated in the database of all GO IDs; BgRatio: The ratio of the number of genes in the GO database annotated by the GO ID and the number of genes in the GO database annotated by any GO ID; pvalue: Unadjusted p-value; p.adjust: FDR p-value using the Benjamini-Hochberg method; qvalue: Alternative method of FDR correction; geneID - List of genes in the query set that are annotated by the GO ID; count: Number of genes in the query set that are annotated by the GO ID

**Supplemental Table 3.** Genes Differentially Expressed in Both Female versus Male Hofbauer Cells and Microglia

Footer: Mg: microglia; HBC: Hofbauer cell; padj: Benjamini Hochberg adjusted p-values. Direction of dysregulation is in females relative to males, so negative log2fold change values indicate a downregulation in females relative to males, and positive log2fold change upregulation.

**Supplemental Table 4.** Working files for the Ingenuity Pathways (IPA) analyses

**Supplemental Table 5.** IPA Analyses for Female versus Male Microglia (Tab 1, Canonical Pathways; Tab 2, Upstream Regulators; Tab 3, Downstream Effects) Tab 1 Footer: Significantly Dysregulated Canonical Pathways in Female versus Male Microglia. Significantly dysregulated pathways defined as those with BH p-value < 0.05 and/or absolute Z-score ≥ 2; -log (BH p-value) > 1.3 corresponds to BH p-value < 0.05; Ratio: number of dysregulated genes in the gene set in the canonical pathway/total genes in the canonical pathway; Differentially-expressed genes in this pathway: genes in our DEG gene set that are component genes of the pathway. Directionality is for female versus male, so upregulated/activated is activated in females.

Tab 2 Footer: Significantly Dysregulated Upstream Regulators in Female versus Male Microglia. Significant dysregulation of Upstream Regulators defined as BH p-value < 0.01 (given large number of upstream regulators with BH p-value < 0.05) and/or absolute Z-score ≥ 2, and involving at least 3 genes in the dataset. Activation is predicted by IPA if Z-score ≥ 2, Inhibition predicted if Z-score ≤ −2.0. Directionality is for female versus male, so upregulated/activated is activated in females. Tab 3 Footer: Significantly Dysregulated Molecular and Cellular Functions and Physiological Systems in Female versus Male Microglia. Significantly dysregulated functions defined by BH p-value < 0.05 and/or absolute Z-score ≥ 2, and involving at least 3 genes in the dataset. Predicted activation is “increased” if Z-score ≥ 2, “decreased” if Z-score ≤ −2.0. Directionality is for female versus male, so increased activation is increased in females.

**Supplemental Table 6.** IPA Analyses for Female versus Male Hofbauer Cells (Tab 1, Upstream Regulators; Tab 2 Downstream Effects)

Tab 1 Footer: Significantly Dysregulated Upstream Regulators in Female versus Male Hofbauer Cells. Significance defined as BH-p < 0.01 and/or absolute Z-score ≥ 2, and involving at least 3 genes in the dataset. Directionality is for female versus male, so upregulated/activated is activated in females.

Tab 2 Footer: Significantly Dysregulated Molecular and Cellular Functions and Physiological Systems in Female versus Male Hofbauer Cells. Significantly dysregulated functions defined by BH p-value < 0.05 and/or absolute Z-score ≥ 2, and involving at least 3 genes in the dataset. Directionality is for female versus male, so upregulated/activated is activated in females.

**Supplemental Table 7.** IPA Analyses for PAMMs from Female versus Male Pregnancies (Tab 1, Upstream Regulators; Tab 2, Downstream Effects)

Tab 1 Footer: Significantly Dysregulated Upstream Regulators in Female vs Male PAMMs. Significant dysregulation of Upstream Regulators defined as BH p-value < 0.01 (given large number of upstream regulators with BH p-value < 0.05) and/or absolute Zscore≥2, and involving at least 3 genes in the dataset. Activation is predicted by IPA if Z-score≥2, Inhibition predicted if Z-score ≤-2.0. Directionality is for female versus male, so upregulated is upregulated in females.

Tab 2 Footer: Significantly Dysregulated Molecular and Cellular Functions and Physiological Systems in Female versus Male PAMMs. Significantly dysregulated functions defined by BH p-value < 0.05 and/or absolute Z-score ≥2, and involving at least 3 genes in the dataset.

## Author Contributions

A.M.C. and R.B. contributed equally, and as co-first authors, may re-arrange their names to appear first when referring to this publication (e.g. on a CV). A.G.E. and S.D.B. conceived the study and, together with A.M.C. and R.B., designed the experiments. Acquisition of data: A.M.C., R.B., S.K., E.A.B., L.L.S., A.G.E. Analysis and interpretation of data: R.B., A.M.C., B.A.D, D.K.S., A.G.E. Drafting of the manuscript: A.M.C, R.B. (bioinformatics methodology), A.G.E. Revising the manuscript critically for important intellectual content: A.M.C., R.B., L.L.S., S.K., E.A.B., B.A.D., R.H.P., D.K.S., S.D.B., A.G.E. All authors have given final approval for submission.

## Acknowledgements

We thank Dr. Abigail Groff and Dr. David Page for advice regarding identification of male versus female macrophages.

## References

Abels, E.R., Nieland, L., Hickman, S., Broekman, M.L.D., El Khoury, J., and Maas, S.L.N. (2021). Comparative Analysis Identifies Similarities between the Human and Murine Microglial Sensomes. Int J Mol Sci 22, 1495.

Ajami, B., Bennett, J.L., Krieger, C., Tetzlaff, W., and Rossi, F.M.V. (2007). Local self-renewal can sustain CNS microglia maintenance and function throughout adult life. Nat Neurosci 10, 1538–1543.

Aran, D., Looney, A.P., Liu, L., Wu, E., Fong, V., Hsu, A., Chak, S., Naikawadi, R.P., Wolters, P.J., Abate, A.R., et al. (2019). Reference-based analysis of lung single-cell sequencing reveals a transitional profibrotic macrophage. Nat Immunol 20, 163–172.

Ayata, P., Badimon, A., Strasburger, H.J., Duff, M.K., Montgomery, S.E., Loh, Y.-H.E., Ebert, A., Pimenova, A.A., Ramirez, B.R., Chan, A.T., et al. (2018). Epigenetic regulation of brain region-specific microglia clearance activity. Nat Neurosci 21, 1049–1060.

Bachiller, S., Jiménez-Ferrer, I., Paulus, A., Yang, Y., Swanberg, M., Deierborg, T., and Boza-Serrano, A. (2018). Microglia in Neurological Diseases: A Road Map to Brain-Disease Dependent-Inflammatory Response. Frontiers in Cellular Neuroscience 12, 488.

Bian, Z., Gong, Y., Huang, T., Lee, C.Z.W., Bian, L., Bai, Z., Shi, H., Zeng, Y., Liu, C., He, J., et al. (2020). Deciphering human macrophage development at single-cell resolution. Nature 582, 571–576.

Bilbo, S.D., and Tsang, V. (2010). Enduring consequences of maternal obesity for brain inflammation and behavior of offspring. The FASEB Journal 24, 2104–2115.

Bilbo, S.D., Biedenkapp, J.C., Der-Avakian, A., Watkins, L.R., Rudy, J.W., and Maier, S.F. (2005). Neonatal infection-induced memory impairment after lipopolysaccharide in adulthood is prevented via caspase-1 inhibition. J Neurosci 25, 8000–8009.

Bilimoria, P.M., and Stevens, B. (2015). Microglia function during brain development: New insights from animal models. Brain Res 1617, 7–17.

Bordeleau, M., Carrier, M., Luheshi, G.N., and Tremblay, M.-È. (2019). Microglia along sex lines: From brain colonization, maturation and function, to implication in neurodevelopmental disorders. Semin Cell Dev Biol 94, 152–163.

Bordt, E.A., Block, C.L.,. Petrozziello, T.,. Sadri-Vakili, G.,. Smith, C.J.,. Edlow, A.G.,. Bilbo, S.D. (2020). Isolation of Microglia from Mouse or Human Tissue. STAR Protocols.

Bordt, E.A., Shook, L.L., Atyeo, C., Pullen, K.M., Guzman, R.M.D., Meinsohn, M.-C., Chauvin, M., Fischinger, S., Yockey, L.J., James, K., et al. (2021). Sexually dimorphic placental responses to maternal SARS-CoV-2 infection.

Bruttger, J., Karram, K., Wörtge, S., Regen, T., Marini, F., Hoppmann, N., Klein, M., Blank, T., Yona, S., Wolf, Y., et al. (2015). Genetic Cell Ablation Reveals Clusters of Local Self-Renewing Microglia in the Mammalian Central Nervous System. Immunity 43, 92–106.

Burton, G.J., Woods, A.W., Jauniaux, E., and Kingdom, J.C.P. (2009). Rheological and Physiological Consequences of Conversion of the Maternal Spiral Arteries for Uteroplacental Blood Flow during Human Pregnancy. Placenta 30, 473–482.

Butler, A., Hoffman, P., Smibert, P., Papalexi, E., and Satija, R. (2018). Integrating single-cell transcriptomic data across different conditions, technologies, and species. Nat Biotechnol 36, 411–420.

Ceasrine, A.M., Devlin, B.A., Bolton, J.L., Jo, Y.C., Huynh, C., Patrick, B., Washington, K., Joo, F., CamposSalazar, A.B., Lockshin, E.R., et al. (2021). Maternal diet disrupts the placenta-brain axis in a sex-specific manner.

Chang, M.D., Pollard, J.W., Khalili, H., Goyert, S.M., and Diamond, B. (1993). Mouse placental macrophages have a decreased ability to present antigen. Proceedings of the National Academy of Sciences 90, 462–466.

Chen, C.-Y., Liu, S.-H., Chen, C.-Y., Chen, P.-C., and Chen, C.-P. (2015a). Human placenta-derived multipotent mesenchymal stromal cells involved in placental angiogenesis via the PDGF-BB and STAT3 pathways. Biol Reprod 93, 103.

Chen, M.-T., Dong, L., Zhang, X.-H., Yin, X.-L., Ning, H.-M., Shen, C., Su, R., Li, F., Song, L., Ma, Y.-N., et al. (2015b). ZFP36L1 promotes monocyte/macrophage differentiation by repressing CDK6. Sci Rep 5, 16229.

Chen, S., Zhao, S., Dalman, C., Karlsson, H., and Gardner, R. (2021). Association of maternal diabetes with neurodevelopmental disorders: autism spectrum disorders, attention-deficit/hyperactivity disorder and intellectual disability. International Journal of Epidemiology 50, 459–474.

Cindrova-Davies, T., Jauniaux, E., Elliot, M.G., Gong, S., Burton, G.J., and Charnock-Jones, D.S. (2017). RNA-seq reveals conservation of function among the yolk sacs of human, mouse, and chicken. PNAS 114, E4753–E4761.

Clifton, V.L., and Murphy, V.E. (2004). Maternal asthma as a model for examining fetal sex-specific effects on maternal physiology and placental mechanisms that regulate human fetal growth. Placenta 25 Suppl A, S45–52.

Cvitic, S., Longtine, M.S., Hackl, H., Wagner, K., Nelson, M.D., Desoye, G., and Hiden, U. (2013). The human placental sexome differs between trophoblast epithelium and villous vessel endothelium. PLoS One 8, e79233.

Edlow, A.G. (2017). Maternal obesity and neurodevelopmental and psychiatric disorders in offspring. Prenat Diagn 37, 95–110.

Edlow, A., Hui, L., Wick, H., Fried, I., and Bianchi, D. (2016a). Assessing the fetal effects of maternal obesity via transcriptomic analysis of cord blood: a prospective case–control study. BJOG: An International Journal of Obstetrics & Gynaecology 123, 180–189.

Edlow, A.G., Slonim, D.K., Wick, H.C., Hui, L., and Bianchi, D.W. (2015). The pathway not taken: understanding ‘omics data in the perinatal context. American Journal of Obstetrics & Gynecology 213, 59.e1–59.e172.

Edlow, A.G., Guedj, F., Pennings, J.L.A., Sverdlov, D., Neri, C., and Bianchi, D.W. (2016b). Males are from Mars, and females are from Venus: sex-specific fetal brain gene expression signatures in a mouse model of maternal diet-induced obesity. American Journal of Obstetrics and Gynecology 214, 623.e1–623.e10.

Edlow, A.G., Glass, R.M., Smith, C.J., Tran, P.K., James, K., and Bilbo, S. (2018). Placental Macrophages: A Window Into Fetal Microglial Function in Maternal Obesity. Int J Dev Neurosci.

Edlow, A.G., Guedj, F., Sverdlov, D., Pennings, J.L.A., and Bianchi, D.W. (2019). Significant Effects of Maternal Diet During Pregnancy on the Murine Fetal Brain Transcriptome and Offspring Behavior. Front Neurosci 13, 1335.

Frick, L., and Pittenger, C. (2016). Microglial Dysregulation in OCD, Tourette Syndrome, and PANDAS. J Immunol Res 2016, 8606057.

Gekas, C., Rhodes, K.E., Van Handel, B., Chhabra, A., Ueno, M., and Mikkola, H.K.A. (2010). Hematopoietic stem cell development in the placenta. Int J Dev Biol 54, 1089–1098.

Ginhoux, F., and Prinz, M. (2015). Origin of microglia: current concepts and past controversies. Cold Spring Harb Perspect Biol 7, a020537.

Ginhoux, F., Greter, M., Leboeuf, M., Nandi, S., See, P., Gokhan, S., Mehler, M.F., Conway, S.J., Ng, L.G., Stanley, E.R., et al. (2010). Fate Mapping Analysis Reveals That Adult Microglia Derive from Primitive Macrophages. Science 330, 841–845.

Godin, I., and Cumano, A. (2002). The hare and the tortoise: an embryonic haematopoietic race. Nat Rev Immunol 2, 593–604.

Gomez Perdiguero, E., Schulz, C., and Geissmann, F. (2013). Development and homeostasis of “resident” myeloid cells: the case of the microglia. Glia 61, 112–120.

Gomez Perdiguero, E., Klapproth, K., Schulz, C., Busch, K., Azzoni, E., Crozet, L., Garner, H., Trouillet, C., de Bruijn, M.F., Geissmann, F., et al. (2015). Tissue-resident macrophages originate from yolk-sacderived erythro-myeloid progenitors. Nature 518, 547–551.

Gonzalez, T.L., Sun, T., Koeppel, A.F., Lee, B., Wang, E.T., Farber, C.R., Rich, S.S., Sundheimer, L.W., Buttle, R.A., Chen, Y.-D.I., et al. (2018). Sex differences in the late first trimester human placenta transcriptome. Biol Sex Differ 9, 4.

Greter, M., Lelios, I., and Croxford, A.L. (2015). Microglia Versus Myeloid Cell Nomenclature during Brain Inflammation. Frontiers in Immunology 6, 249.

Hafemeister, C., and Satija, R. (2019). Normalization and variance stabilization of single-cell RNA-seq data using regularized negative binomial regression. Genome Biology 20, 296.

Haimon, Z., Volaski, A., Orthgiess, J., Boura-Halfon, S., Varol, D., Shemer, A., Yona, S., Zuckerman, B., David, E., Chappell-Maor, L., et al. (2018). Re-evaluating microglia expression profiles using RiboTag and cell isolation strategies. Nat Immunol 19, 636–644.

Haley, M.J., Brough, D., Quintin, J., and Allan, S.M. (2019). Microglial Priming as Trained Immunity in the Brain. Neuroscience 405, 47–54.

Hammond, T.R., Dufort, C., Dissing-Olesen, L., Giera, S., Young, A., Wysoker, A., Walker, A.J., Gergits, F., Segel, M., Nemesh, J., et al. (2019). Single-Cell RNA Sequencing of Microglia throughout the Mouse Lifespan and in the Injured Brain Reveals Complex Cell-State Changes. Immunity 50, 253–271.e6.

Hanamsagar, R., and Bilbo, S.D. (2016). Sex differences in neurodevelopmental and neurodegenerative disorders: Focus on microglial function and neuroinflammation during development. J Steroid Biochem Mol Biol 160, 127–133.

Hanamsagar, R., and Bilbo, S.D. (2017). Environment Matters: Microglia Function and Dysfunction in a Changing World. Curr Opin Neurobiol 47, 146–155.

Hao, Y., Hao, S., Andersen-Nissen, E., Mauck, W.M., Zheng, S., Butler, A., Lee, M.J., Wilk, A.J., Darby, C., Zager, M., et al. (2021). Integrated analysis of multimodal single-cell data. Cell 184, 3573–3587.e29.

Hemberger, M., Hanna, C.W., and Dean, W. (2020). Mechanisms of early placental development in mouse and humans. Nat Rev Genet 21, 27–43.

Imakawa, K., Dhakal, P., Kubota, K., Kusama, K., Chakraborty, D., Karim Rumi, M.A., and Soares, M.J. (2016). CITED2 modulation of trophoblast cell differentiation: insights from global transcriptome analysis. Reproduction 151, 509–516.

Ingenuity Systems (2021a). Calculating and Interpreting the p-values for Functions, Pathways and Lists in IPA.

Ingenuity Systems (2021b). Ingenuity Upstream Regulator Analysis in IPA.

Ji, Y.Q., Zhang, Y.Q., Li, M.Q., Du, M.R., Wei, W.W., and Li, D.J. (2011). EPO improves the proliferation and inhibits apoptosis of trophoblast and decidual stromal cells through activating STAT-5 and inactivating p38 signal in human early pregnancy. Int J Clin Exp Pathol 4, 765–774.

Johnson, E.L., and Chakraborty, R. (2012). Placental Hofbauer cells limit HIV-1 replication and potentially offset mother to child transmission (MTCT) by induction of immunoregulatory cytokines. Retrovirology 9, 101.

Johnson, E.L., Boggavarapu, S., Johnson, E.S., Lal, A.A., Agrawal, P., Bhaumik, S.K., Murali-Krishna, K., and Chakraborty, R. (2018). Human Cytomegalovirus Enhances Placental Susceptibility and Replication of Human Immunodeficiency Virus Type 1 (HIV-1), Which May Facilitate In Utero HIV-1 Transmission. J Infect Dis 218, 1464–1473.

Jurado, K.A., Simoni, M.K., Tang, Z., Uraki, R., Hwang, J., Householder, S., Wu, M., Lindenbach, B.D., Abrahams, V.M., Guller, S., et al. Zika virus productively infects primary human placenta-specific macrophages. JCI Insight 1, e88461.

Khalili, H., Deshpande, R., and Chang, M.Y. (1997). The defective antigen-presenting activity of murine fetal macrophage cell lines. Immunology 92, 487–493.

Kieffer, T.E.C., Laskewitz, A., Faas, M.M., Scherjon, S.A., Erwich, J.J.H.M., Gordijn, S.J., and Prins, J.R. (2018). Lower FOXP3 mRNA Expression in First-Trimester Decidual Tissue from Uncomplicated Term Pregnancies with a Male Fetus. J Immunol Res 2018, 1950879.

Kipkeew, F., Kirsch, M., Klein, D., Wuelling, M., Winterhager, E., and Gellhaus, A. (2016). CCN1 (CYR61) and CCN3 (NOV) signaling drives human trophoblast cells into senescence and stimulates migration properties. Cell Adh Migr 10, 163–178.

Kracht, L., Borggrewe, M., Eskandar, S., Brouwer, N., Chuva de Sousa Lopes, S.M., Laman, J.D., Scherjon, S.A., Prins, J.R., Kooistra, S.M., and Eggen, B.J.L. (2020). Human fetal microglia acquire homeostatic immune-sensing properties early in development. Science 369, 530–537.

Krämer, A., Green, J., Pollard, J., and Tugendreich, S. (2014). Causal analysis approaches in Ingenuity Pathway Analysis. Bioinformatics 30, 523–530.

Lenz, K.M., and McCarthy, M.M. (2015). A Starring Role for Microglia in Brain Sex Differences. Neuroscientist 21, 306–321.

Li, J.-W., Zong, Y., Cao, X.-P., Tan, L., and Tan, L. (2018). Microglial priming in Alzheimer’s disease. Ann Transl Med 6, 176.

Liu, Y., Fan, X., Wang, R., Lu, X., Dang, Y.-L., Wang, H., Lin, H.-Y., Zhu, C., Ge, H., Cross, J.C., et al. (2018). Single-cell RNA-seq reveals the diversity of trophoblast subtypes and patterns of differentiation in the human placenta. Cell Res 28, 819–832.

Love, M.I., Huber, W., and Anders, S. (2014). Moderated estimation of fold change and dispersion for RNA-seq data with DESeq2. Genome Biology 15, 550.

Lu-Culligan, A., Chavan, A.R., Vijayakumar, P., Irshaid, L., Courchaine, E.M., Milano, K.M., Tang, Z., Pope, S.D., Song, E., Vogels, C.B.F., et al. (2021). SARS-CoV-2 infection in pregnancy is associated with robust inflammatory response at the maternal-fetal interface. MedRxiv 2021.01.25.21250452.

Maciejewski, J.P., Bruening, E.E., Donahue, R.E., Sellers, S.E., Carter, C., Young, N.S., and St Jeor, S. (1993). Infection of mononucleated phagocytes with human cytomegalovirus. Virology 195, 327–336.

Madisen, L., Zwingman, T.A., Sunkin, S.M., Oh, S.W., Zariwala, H.A., Gu, H., Ng, L.L., Palmiter, R.D., Hawrylycz, M.J., Jones, A.R., et al. (2010). A robust and high-throughput Cre reporting and characterization system for the whole mouse brain. Nat Neurosci 13, 133–140.

Marschallinger, J., Iram, T., Zardeneta, M., Lee, S.E., Lehallier, B., Haney, M.S., Pluvinage, J.V., Mathur, V., Hahn, O., Morgens, D.W., et al. (2020). Lipid-droplet-accumulating microglia represent a dysfunctional and proinflammatory state in the aging brain. Nat Neurosci 23, 194–208.

Masuda, T., Sankowski, R., Staszewski, O., Böttcher, C., Amann, L., Sagar, Scheiwe, C., Nessler, S., Kunz, P., van Loo, G., et al. (2019). Spatial and temporal heterogeneity of mouse and human microglia at single-cell resolution. Nature 566, 388–392.

Matcovitch-Natan, O., Winter, D.R., Giladi, A., Vargas Aguilar, S., Spinrad, A., Sarrazin, S., Ben-Yehuda, H., David, E., Zelada Gonzalez, F., Perrin, P., et al. (2016). Microglia development follows a stepwise program to regulate brain homeostasis. Science 353, aad8670–aad8670.

McGinnis, C.S., Murrow, L.M., and Gartner, Z.J. (2019). DoubletFinder: Doublet Detection in Single-Cell RNA Sequencing Data Using Artificial Nearest Neighbors. Cels 8, 329–337.e4.

Megli, C., and Coyne, C.B. (2020). Gatekeepers of the fetus: Characterization of placental macrophages. Journal of Experimental Medicine 218.

Mestas, J., and Hughes, C.C.W. (2004). Of Mice and Not Men: Differences between Mouse and Human Immunology. The Journal of Immunology 172, 2731–2738.

Mezouar, S., Katsogiannou, M., Ben Amara, A., Bretelle, F., and Mege, J.-L. (2021). Placental macrophages: Origin, heterogeneity, function and role in pregnancy-associated infections. Placenta 103, 94–103.

Migliaccio, G., Migliaccio, A.R., Petti, S., Mavilio, F., Russo, G., Lazzaro, D., Testa, U., Marinucci, M., and Peschle, C. (1986). Human embryonic hemopoiesis. Kinetics of progenitors and precursors underlying the yolk sac----liver transition. J Clin Invest 78, 51–60.

Moffett, A., and Loke, C. (2006). Immunology of placentation in eutherian mammals. Nat Rev Immunol 6, 584–594.

Mor, G., and Abrahams, V.M. (2003). Potential role of macrophages as immunoregulators of pregnancy. Reprod Biol Endocrinol 1, 119.

Na, Q., Chudnovets, A., Liu, J., Lee, J.Y., Dong, J., Shin, N., Elsayed, N., Lei, J., and Burd, I. (2021). Placental Macrophages Demonstrate Sex-Specific Response to Intrauterine Inflammation and May Serve as a Marker of Perinatal Neuroinflammation. Journal of Reproductive Immunology 147, 103360.

Nogueira Avelar E Silva, R., Yu, Y., Liew, Z., Vested, A., Sørensen, H.T., and Li, J. (2021). Associations of Maternal Diabetes During Pregnancy With Psychiatric Disorders in Offspring During the First 4 Decades of Life in a Population-Based Danish Birth Cohort. JAMA Netw Open 4, e2128005.

Noronha, L. de, Zanluca, C., Azevedo, M.L.V., Luz, K.G., and Santos, C.N.D.D. (2016). Zika virus damages the human placental barrier and presents marked fetal neurotropism. Mem Inst Oswaldo Cruz 111, 287–293.

Palis, J., and Yoder, M.C. (2001). Yolk-sac hematopoiesis: the first blood cells of mouse and man. Exp Hematol 29, 927–936.

Paolicelli, R.C., Bolasco, G., Pagani, F., Maggi, L., Scianni, M., Panzanelli, P., Giustetto, M., Ferreira, T.A., Guiducci, E., Dumas, L., et al. (2011). Synaptic pruning by microglia is necessary for normal brain development. Science 333, 1456–1458.

Pinhal-Enfield, G., Vasan, N.S., and Leibovich, S.J. (2012). The Role of Macrophages in the Placenta (IntechOpen).

PrabhuDas, M., Bonney, E., Caron, K., Dey, S., Erlebacher, A., Fazleabas, A., Fisher, S., Golos, T., Matzuk, M., McCune, J.M., et al. (2015). Immune mechanisms at the maternal-fetal interface: perspectives and challenges. Nat Immunol 16, 328–334.

Quicke, K.M., Bowen, J.R., Johnson, E.L., McDonald, C.E., Ma, H., O’Neal, J.T., Rajakumar, A., Wrammert, J., Rimawi, B.H., Pulendran, B., et al. (2016). Zika Virus Infects Human Placental Macrophages. Cell Host Microbe 20, 83–90.

Reyes, L., and Golos, T.G. (2018). Hofbauer Cells: Their Role in Healthy and Complicated Pregnancy. Frontiers in Immunology 9.

Rhodes, K.E., Gekas, C., Wang, Y., Lux, C.T., Francis, C.S., Chan, D.N., Conway, S., Orkin, S.H., Yoder, M.C., and Mikkola, H.K.A. (2008). The Emergence of Hematopoietic Stem Cells is Initiated in the Placental Vasculature in the Absence of Circulation. Cell Stem Cell 2, 252–263.

Rosenberg, A.Z., Yu, W., Hill, D.A., Reyes, C.A., and Schwartz, D.A. (2017). Placental Pathology of Zika Virus: Viral Infection of the Placenta Induces Villous Stromal Macrophage (Hofbauer Cell) Proliferation and Hyperplasia. Arch Pathol Lab Med 141, 43–48.

Rosenfeld, C.S. (2015). Sex-Specific Placental Responses in Fetal Development. Endocrinology 156, 3422–3434.

Salter, M.W., and Stevens, B. (2017). Microglia emerge as central players in brain disease. Nat Med 23, 1018–1027.

Sasmono, R.T., Oceandy, D., Pollard, J.W., Tong, W., Pavli, P., Wainwright, B.J., Ostrowski, M.C., Himes, S.R., and Hume, D.A. (2003). A macrophage colony-stimulating factor receptor–green fluorescent protein transgene is expressed throughout the mononuclear phagocyte system of the mouse. Blood 101, 1155–1163.

Satija, R., Farrell, J.A., Gennert, D., Schier, A.F., and Regev, A. (2015). Spatial reconstruction of single-cell gene expression data. Nat Biotechnol 33, 495–502.

Schafer, D.P., Lehrman, E.K., Kautzman, A.G., Koyama, R., Mardinly, A.R., Yamasaki, R., Ransohoff, R.M., Greenberg, M.E., Barres, B.A., and Stevens, B. (2012). Microglia sculpt postnatal neural circuits in an activity and complement-dependent manner. Neuron 74, 691–705.

Schindelin, J., Arganda-Carreras, I., Frise, E., Kaynig, V., Longair, M., Pietzsch, T., Preibisch, S., Rueden, C., Saalfeld, S., Schmid, B., et al. (2012). Fiji: an open-source platform for biological-image analysis. Nat Methods 9, 676–682.

Scott, N.M., Hodyl, N.A., Murphy, V.E., Osei-Kumah, A., Wyper, H., Hodgson, D.M., Smith, R., and Clifton, V.L. (2009). Placental cytokine expression covaries with maternal asthma severity and fetal sex. J Immunol 182, 1411–1420.

Semmes, E.C., and Coyne, C.B. (2022). Innate immune defenses at the maternal-fetal interface. Current Opinion in Immunology 74, 60–67.

Shook, L.L., Kislal, S., and Edlow, A.G. (2020). Fetal brain and placental programming in maternal obesity: A review of human and animal model studies. Prenat Diagn 40, 1126–1137.

Sierra, A., Encinas, J.M., Deudero, J.J.P., Chancey, J.H., Enikolopov, G., Overstreet-Wadiche, L.S., Tsirka, S.E., and Maletic-Savatic, M. (2010). Microglia shape adult hippocampal neurogenesis through apoptosis-coupled phagocytosis. Cell Stem Cell 7, 483–495.

Simoni, M.K., Jurado, K.A., Abrahams, V.M., Fikrig, E., and Guller, S. (2017). Zika Virus Infection of Hofbauer Cells. Am J Reprod Immunol 77, 10.1111/aji.12613.

Soneson, C., and Robinson, M.D. (2018). Bias, robustness and scalability in single-cell differential expression analysis. Nat Methods 15, 255–261.

Squair, J.W., Gautier, M., Kathe, C., Anderson, M.A., James, N.D., Hutson, T.H., Hudelle, R., Qaiser, T., Matson, K.J.E., Barraud, Q., et al. (2021). Confronting false discoveries in single-cell differential expression. Nat Commun 12, 5692.

Stremmel, C., Schuchert, R., Wagner, F., Thaler, R., Weinberger, T., Pick, R., Mass, E., IshikawaAnkerhold, H.C., Margraf, A., Hutter, S., et al. (2018). Yolk sac macrophage progenitors traffic to the embryo during defined stages of development. Nat Commun 9, 75.

Stuart, T., Butler, A., Hoffman, P., Hafemeister, C., Papalexi, E., Mauck, W.M., Hao, Y., Stoeckius, M., Smibert, P., and Satija, R. (2019). Comprehensive Integration of Single-Cell Data. Cell 177, 18881902.e21.

Stumpo, D.J., Byrd, N.A., Phillips, R.S., Ghosh, S., Maronpot, R.R., Castranio, T., Meyers, E.N., Mishina, Y., and Blackshear, P.J. (2004). Chorioallantoic fusion defects and embryonic lethality resulting from disruption of Zfp36L1, a gene encoding a CCCH tandem zinc finger protein of the Tristetraprolin family. Mol Cell Biol 24, 6445–6455.

Sun, T., Gonzalez, T.L., Deng, N., DiPentino, R., Clark, E.L., Lee, B., Tang, J., Wang, Y., Stripp, B.R., Yao, C., et al. (2020). Sexually Dimorphic Crosstalk at the Maternal-Fetal Interface. J Clin Endocrinol Metab 105, dgaa503.

Suryawanshi, H., Morozov, P., Straus, A., Sahasrabudhe, N., Max, K.E.A., Garzia, A., Kustagi, M., Tuschl, T., and Williams, Z. (2018). A single-cell survey of the human first-trimester placenta and decidua. Sci Adv 4, eaau4788.

Takahashi, K., Naito, M., Katabuchi, H., and Higashi, K. (1991). Development, Differentiation, and Maturation of Macrophages in the Chorionic Villi of Mouse Placenta With Special Reference to the Origin of Hofbauer Cells. J Leukoc Biol 50, 57–68.

Takashina, T. (1987). Haemopoiesis in the human yolk sac. J Anat 151, 125–135.

Tay, T.L., Béchade, C., D’Andrea, I., St-Pierre, M.-K., Henry, M.S., Roumier, A., and Tremblay, M.-E. (2018). Microglia Gone Rogue: Impacts on Psychiatric Disorders across the Lifespan. Frontiers in Molecular Neuroscience 10, 421.

Thomas, J.R., Appios, A., Zhao, X., Dutkiewicz, R., Donde, M., Lee, C.Y.C., Naidu, P., Lee, C., Cerveira, J., Liu, B., et al. (2020). Phenotypic and functional characterization of first-trimester human placental macrophages, Hofbauer cells. J Exp Med 218, e20200891.

Tsang, J.C.H., Vong, J.S.L., Ji, L., Poon, L.C.Y., Jiang, P., Lui, K.O., Ni, Y.-B., To, K.F., Cheng, Y.K.Y., Chiu, R.W.K., et al. (2017). Integrative single-cell and cell-free plasma RNA transcriptomics elucidates placental cellular dynamics. PNAS 114, E7786–E7795.

Van Lieshout, R.J., and Voruganti, L.P. (2008). Diabetes mellitus during pregnancy and increased risk of schizophrenia in offspring: a review of the evidence and putative mechanisms. J Psychiatry Neurosci 33, 395–404.

Vento-Tormo, R., Efremova, M., Botting, R.A., Turco, M.Y., Vento-Tormo, M., Meyer, K.B., Park, J.-E., Stephenson, E., Polański, K., Goncalves, A., et al. (2018). Single-cell reconstruction of the early maternal– fetal interface in humans. Nature 563, 347–353.

Wickham, H., Averick, M., Bryan, J., Chang, W., McGowan, L.D., François, R., Grolemund, G., Hayes, A., Henry, L., Hester, J., et al. (2019). Welcome to the Tidyverse. Journal of Open Source Software 4, 1686.

Williamson, L.L., Sholar, P.W., Mistry, R.S., Smith, S.H., and Bilbo, S.D. (2011). Microglia and Memory: Modulation by Early-Life Infection. J. Neurosci.

Wu, T., Hu, E., Xu, S., Chen, M., Guo, P., Dai, Z., Feng, T., Zhou, L., Tang, W., Zhan, L., et al. (2021). clusterProfiler 4.0: A universal enrichment tool for interpreting omics data. The Innovation 2, 100141.

Yellon, S.M., Greaves, E., Heuerman, A.C., Dobyns, A.E., and Norman, J.E. (2019). Effects of macrophage depletion on characteristics of cervix remodeling and pregnancy in CD11b-dtr mice. Biol Reprod 100, 1386–1394.

Zheng, G.X.Y., Terry, J.M., Belgrader, P., Ryvkin, P., Bent, Z.W., Wilson, R., Ziraldo, S.B., Wheeler, T.D., McDermott, G.P., Zhu, J., et al. (2017). Massively parallel digital transcriptional profiling of single cells. Nat Commun 8, 14049.

Zhou, X., Xu, Y., Ren, S., Liu, D., Yang, N., Han, Q., Kong, S., Wang, H., Deng, W., Qi, H., et al. (2021). Single-cell RNA-seq revealed diverse cell types in the mouse placenta at mid-gestation. Experimental Cell Research 405, 112715.

Zulu, M.Z., Martinez, F.O., Gordon, S., and Gray, C.M. (2019). The Elusive Role of Placental Macrophages: The Hofbauer Cell. JIN 11, 447–456.

