## Supplemental Figures for "Single cell profiling of Hofbauer cells and fetal brain microglia reveals shared programs and functions"

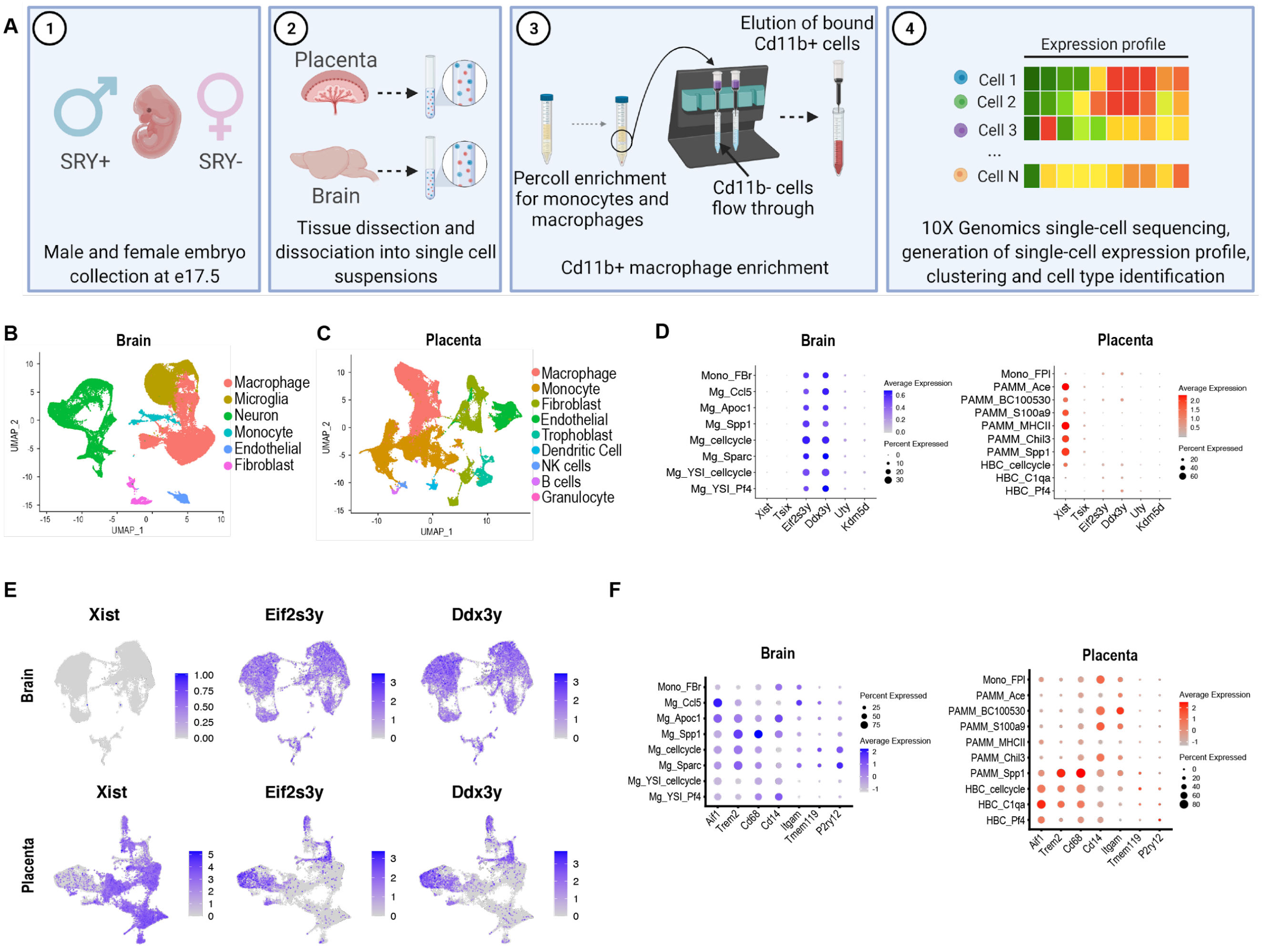


**Supplemental Figure 1: Experimental strategy and identification of fetally- versus maternally-derived macrophage clusters**

**A.** Schematic depicting experimental paradigm. This figure was made using Biorender.

**B.** Uniform manifold Approximation and Projection (UMAP) plots of all brain clusters identified prior to selection for macrophage/monocyte clusters

**C.** UMAP of all placenta clusters identified prior to selection for macrophage/monocyte clusters

**D.** Dot plot of all male-specific marker expression in brain and placenta clusters.

**E.** UMAPs of representative female-specific (*Xist*) and male-specific (*Eif2s3y* and *Ddx3y*) marker expression in brain and placenta clusters from male fetal tissues.

**F.** Dot plot depicting canonical microglia marker expression in brain and placenta clusters

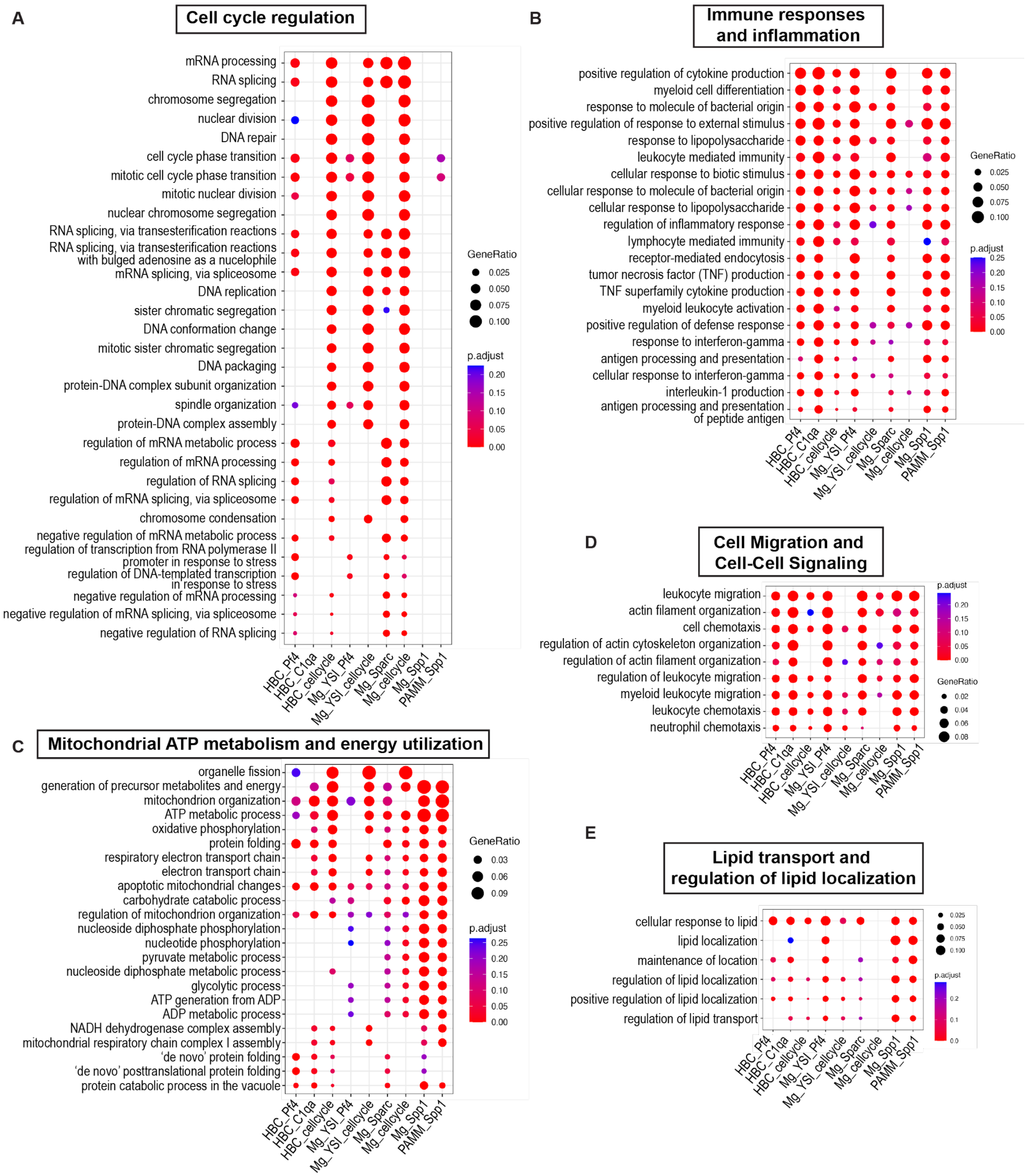


**Supplemental Figure 2: Shared functions between Hofbauer cells and microglia**

**A-E.** Top 15 Gene Ontology (GO) Biological Process enrichment analysis for all Hofbauer cell and microglial clusters, and PAMM_Spp1 cluster, the cluster most similar to Hofbauer cells in its gene expression profile; our clustering suggests PAMM_Spp1 contains maternal cells interacting with the fetal cells in HBC_C1qa. The Top 15 GO functions per cluster have been grouped into broad categories to facilitate understanding of shared common functions that may be performed by Hofbauer cells and microglia. Gene Ratio gives the ratio of the number of genes in the query set that are annotated by the relevant GO category and the number of genes in the query set that are annotated in the database of all GO categories. GO terms with an adjusted p-value < 0.05 were considered significantly enriched.

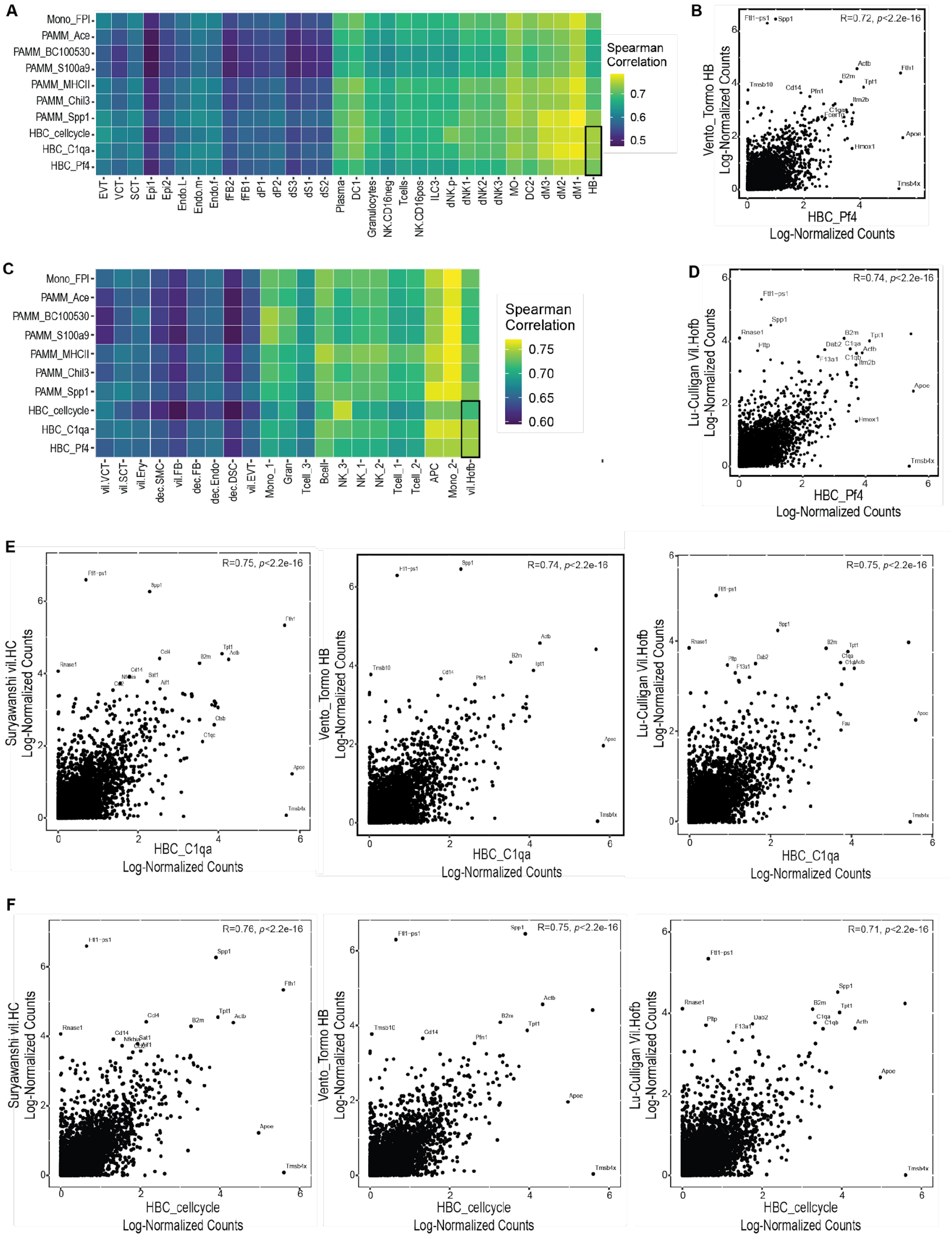

**Supplemental Figure 3: Correlation between murine and human placental macrophage signatures**

**A.** Heatmap displaying Spearman correlation coefficient between our murine single-cell placental clusters and previously published (Vento-Tormo et al., 2018) human placental clusters from first trimester placentas. Hofbauer cell clusters outlined in black. EVT: extravillous trophoblast, VCT: villous cytotrophoblast, SCT: syncytiotrophoblast, Epi1/2: epithelial glandular cells, Endo.L/m/f: endothelial cells (lymphatic/maternal/fetal), fFB1/2: fetal fibroblasts dP1/2: decidual plasma dS1/2/3: decidual stromal cells, DC1/2: dendritic cells, NK.CD16neg: CD16 negative natural killer cells, NK.CD16pos: CD16 positive natural killer cells, ILC3: innate lymphocyte cells, dNK1/2/3: decidual natural killer cells, MO: monocytes dM1/2/3: decidual macrophages, HB: Hofbauer cells.

**B.** Dot plot demonstrating the significant correlation between our Hofbauer cell cluster HBC_Pf4 and the Hofbauer cell cluster identified by Vento-Tormo et al. (2018)

**C.** Heatmap displaying Spearman correlations between our murine single-cell placental clusters and human placental clusters from term placentas (Lu-Culligan et al., 2021). Hofbauer cell clusters outlined in black. vil.VCT: villous cytotrophoblast, vil.SCT:, vil.Ery: villous erythrocyte/blast, dec.SMC: decidual smooth muscle cell, vil.FB: villous fibroblast, dec.Endo: decidual endothelial cell, dec.DSC: decidual stromal cell, vil.EVT: extravillous trophoblast, Mono_1/2: monocyte, Gran: granulocyte, NK_1/2/3: natural killer cell, APC: antigen presenting cell, vil.Hofb: villous Hofbauer cell

**D.** Dot plot demonstrating the significant correlation between our Hofbauer cell cluster HBC_Pf4 and the Hofbauer cell cluster identified by Lu-Culligan et al (2021).

**E.** Dot plots demonstrating significant Spearman's correlations between our Hofbauer cell cluster HBC_C1qa and the Hofbauer cell clusters identified by Suryawanshi, Vento-Tormo, and Lu-Culligan (L to R).

**F.** Dot plots demonstrating significant Spearman's correlations between our Hofbauer cell cluster HBC_cell cycle and the Hofbauer cell clusters identified by Suryawanshi, Vento-Tormo, and Lu-Culligan (L to R).

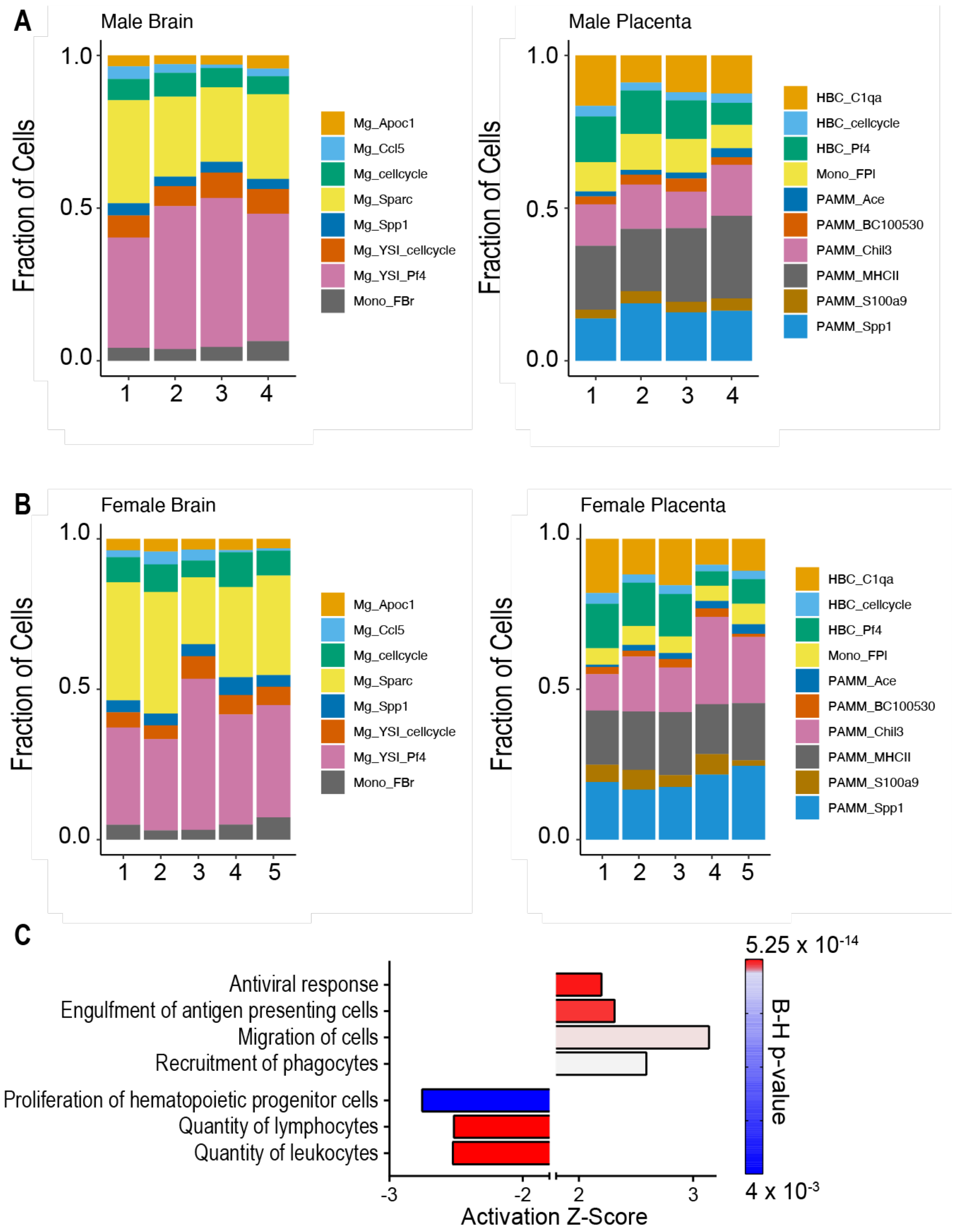


**Supplemental Figure 4: Evaluation of the impact of sex on proportion of cells by cluster and on PAMM molecular and cellular functions.**
**A.** Proportion of cells by cluster in male brain and placenta. X axis depicts male sample numbers (matched fetal brain and placenta), Y axis depicts fraction of cells per cluster.
**B.** Proportion of cells by cluster in female brain and placenta. X axis depicts female sample numbers (matched fetal brain and placenta), Y axis depicts fraction of cells per cluster.
**C.** Up- and downregulated molecular and cellular functions in placenta-associated maternal macrophages and monocytes (PAMMs) from female versus male pregnancy. B-H: Benjamini-Hochberg adjusted p-value
